# DrFARM: Identification and inference for pleiotropic gene in GWAS

**DOI:** 10.1101/2022.11.10.515671

**Authors:** Lap Sum Chan, Gen Li, Eric B. Fauman, Markku Laakso, Michael Boehnke, Peter X.K. Song

## Abstract

In a standard analysis, pleiotropic variants are identified by running separate genome-wide association studies (GWAS) and combining results across traits. But such two-stage statistical approach may lead to spurious results. We propose a new statistical approach, **D**ebiased-**r**egularized **F**actor **A**nalysis **R**egression **M**odel (DrFARM), through a joint regression model for simultaneous analysis of high-dimensional genetic variants and multilevel dependencies. This joint modeling strategy controls overall error to permit universal false discovery rate (FDR) control. DrFARM uses the strengths of the debiasing technique and the Cauchy combination test, both being theoretically justified, to establish a valid post selection inference on pleiotropic variants. Through extensive simulations, we show that DrFARM appropriately controls overall FDR. Applying DrFARM to data on 1,031 metabolites measured on 6,135 men from the Metabolic Syndrome in Men (METSIM) study, we identify 288 new metabolite associations at loci that did not reach statistical significance in prior METSIM metabolite GWAS.

## 1 Introduction

Genetic studies can help identify the contributions of different variants and genes to various processes and pathways. Identifying pleiotropic genes can help us better understand the mechanism of metabolism pathways [1, 2]. Given that technological advances have significantly accelerated the availability of various multi-omics data types (e.g. genomics, epigenomics, transcriptomics, proteomics, metabolomics, glycomics) [3], an unprecedented opportunity arises in the characterization and quantification of pleiotropic genes and genetic variants that regulate multiple phenotypes. However, data analytic techniques to detect pleiotropic genes now lag behind the requirements for increasing high-dimensional data; there are few adequate data analytic methods and software tools available to address the complexity and multimodality of biological data in the detection of pleiotropic genes. Valid statistical methods are essential to explore and understand the underlying biology, generate new hypotheses, and design new experiments to deliver potentially better therapeutics as part of the effort to turn data to knowledge that ultimately improves human quality of life.

Our methods development is largely motivated by the objective of identifying pleiotropic genes for various metabolic traits associated with Type 2 diabetes (T2D) in the **Met**abolic **S**yndrome **i**n **M**en (METSIM) cohort [4], a longitudinal study of 10,197 middle-aged and older Finnish men that seeks to identify genetic variants that contribute to the risk of metabolic and cardiovascular disease. T2D is a complex trait that largely involves the interplay between multiple genes [5, 6]. Discovering pleiotropic genetic variants is one of the key tasks to understand how multiple genetic variants interact in biochemical pathways influencing the risk of developing T2D. Currently, most genome-wide association studies (GWAS) do not formally test for pleiotropy. If testing of pleiotropy is performed, they are based on a single-trait, single-variant analysis approach, which tests for the association of each trait with each variant [7, 8], followed by a second stage of detecting pleiotropic variants using certain GWAS summary statistics [9–12]. However, the linkage disequilibrium (LD) between single nucleotide polymorphisms (SNPs) or variants presents a major challenge in identifying pleiotropic variants. We show that these two-stage approaches that identify genetic pleiotropy based on pairwise marginal association testing cannot control the false discovery rate (FDR) and hence are susceptible to spurious findings.

We introduce DrFARM as a method to identify pleiotropic variants in which confounding by other genetic variants can be adjusted. DrFARM provides a high-dimensional estimation of the coefficients and inference of pleiotropic variants as it is developed to handle data with the number of variants exceeding the sample size. Zhou et al. [13] proposed a sparse multivariate factor analysis regression model (FARM), a high-dimensional joint modeling approach, to detect the so-called “master regulators” (a.k.a. pleiotropic variants), in which they used sparse group lasso regularization [14] to enforce sparsity at both individual-level (entry-level) and group-level (variant-level) [13, 15]. The group sparsity led to the identification of variants being simultaneously associated with multiple traits. The limitation of the sparse multivariate FARM includes that it does not quantify uncertainty and it does not yield FDR control in the discovery of pleiotropic variants. In addition, sparse multivariate FARM ignores relatedness and population structure [16–20].

DrFARM is built upon a post selection debiasing technique to address these limitations, where valid *p*-values are obtained for statistical inference on pleiotropic variants. The debiasing-based post selection (DPS) inference has been studied extensively in the fields of high-dimensional statistics and machine learning [21–24]. This method has seen only limited previous application in genetic data analyses, an area that naturally demands valid DPS inferences [25]. The critical technical challenge in the utility of DPS inferences lies in the estimation of the precision matrix of the predictors, which is the inverse of the covariance matrix of the predictors. This matrix plays a central role in DPS inference as it is used in desparsifying regularized estimates, which are then known to follow asymptotic distributions, and consequently allows for high-dimensional statistical inference, including valid *p*-values generation. Although several methods for precision matrix estimation exist, such as graphical lasso (Glasso) [26], nodewise lasso [21], and quadratic optimization [23], there is no consensus on which method has the best FDR control, sensitivity of parameter tuning, robustness of numerical performance, and computational efficiency. To the best of our knowledge, this paper is the first to conduct a comprehensive comparison of existing precision matrix estimation methods in DPS inference using large-scale simulations, leading to practical guidelines on the use of DPS inference in the analysis of pleiotropic variants. Such knowledge may be applied to many empirical studies with limited sample sizes encountered by other high-dimensional genetic and omics data analyses.

DrFARM: 1) performs a rigorous, valid statistical test via debiasing, to identify potential pleiotropic variants with a proper overall FDR control; 2) accounts for the relatedness and population structure of genetic data in DPS inference; and 3) allows users to choose a precision matrix estimation method in DPS inference. We demonstrate the performance of DrFARM through extensive simulations and make recommendations useful to the application of DrFARM in practical studies. We also reanalyze metabolomics data from the METSIM study to discover new pleiotropic variants and genes.

## 2 Results

### 2.1 Motivating example

We begin with a simple but representative simulation example to motivate the proposed method. We illustrate how pleiotropy may lead to complications in statistical inference. Under the setting of two simulated correlated traits, we illustrate the empirical type I error given by three approaches to identifying pleiotropic variants under the case *P* < *N* : I) the two-stage approach: *p*-values are first obtained using a single-trait, single variant analysis (i.e., univariate *Y*_*j*_, *j* = 1, 2 regressed on single *X*_*i*_, *i* = 1, …, *P*, respectively) and combined for each variant using the Fisher combination test which takes into account the correlation of **Y** = (*Y*_1_, *Y*_2_) [9, 10]; II) MANOVA on multivariate marginal model (i.e., multivariate **Y** regressed on single *X*_*i*_, *i* = 1, …, *P*, respectively); and III) MANOVA on multivariate joint model of *P* variables (i.e., **Y** regressed on **X** = (*X*_1_, …, *X*_*P*_)). Figure 1 shows the average empirical type I error of the three methods. The two methods based on pairwise association testing suffer severely inflated empirical type I error. In particular, the Fisher combination test gets ∼82% average empirical type I error even when the LD between the SNPs was minimal (average *r*^2^ = 0.005 over 1000 replicates). On the other hand, the empirical type I error of the joint MANOVA model is virtually unaffected by the subgroup heterogeneity with a constant ∼5% type I error. This desirable error control is attributed to the fact that the test statistics in the joint modeling adjust for the correlation in traits and SNPs. In contrast, without accounting for the correlation in SNPs, the same MANOVA modeling, when applied to pairwise marginal models, fails to control the overall type I error (∼ 43.5% on average). This simple example implies the need for a joint modeling approach to identifying pleiotropic variants. For illustration, we limited the number of variants equal to that of a set of genomewide significant index variants in the original METSIM marginal analysis as they were the most likely candidates for pleiotropic variants. In practice, it is almost always the case *P > N* (e.g., using 10^−6^ cutoff instead of 5 × 10^−8^). Thus, our development of DrFARM further extends the joint MANOVA modeling approach for the high-dimensional case with *P > N*, which are commonly encountered in the study of pleiotropic variants.

**Fig. 1.**
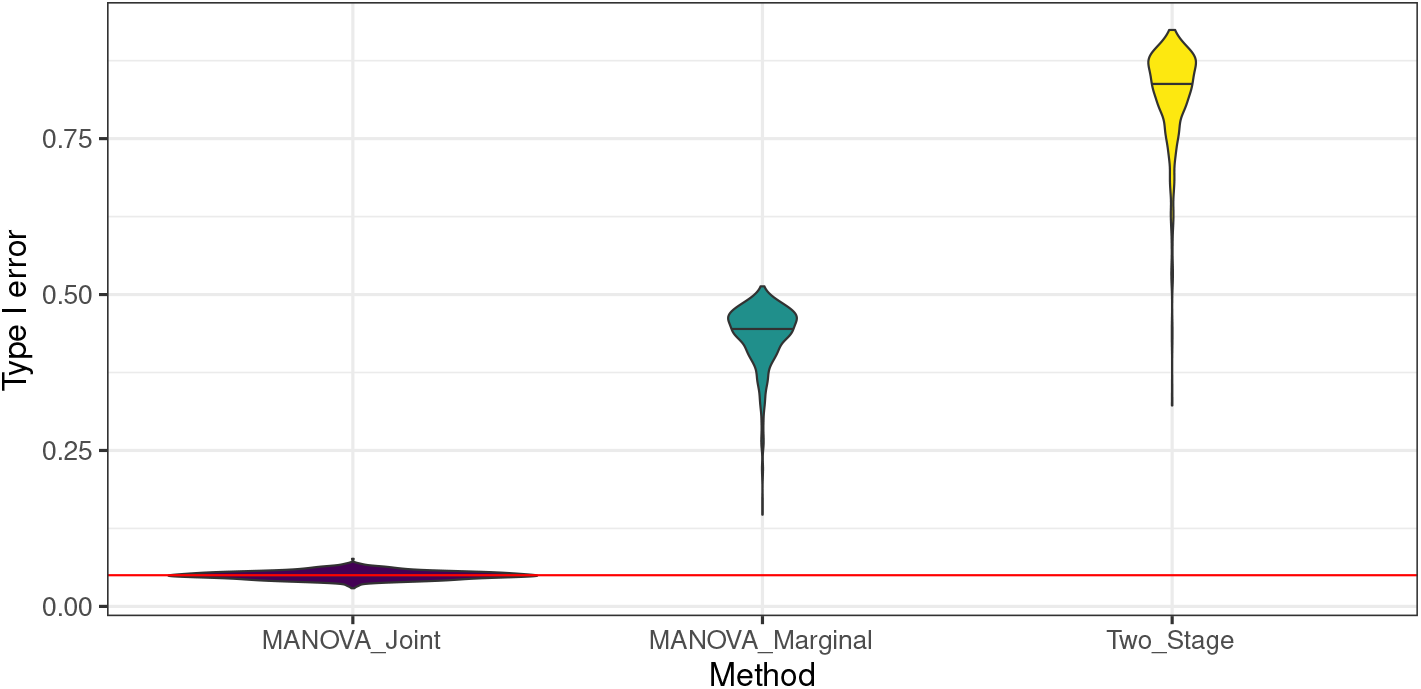
Violin plot of average empirical Type I error for three methods across 1000 replicates: Two stage approach (Two_Stage), marginal MANOVA model (MANOVA_Marginal) and joint MANOVA model (MANOVA_Joint).

### 2.2 Overview

We consider a penalized multivariate regression framework that extends the sparse multivariate FARM [13] (see Section 2 in Methods for more details) to establish valid post selection statistical inference. Compared to traditional linear mixed models in GWAS, DrFARM enables the adjustment for other variants via the high-dimensional joint modeling between *P* variants and *Q* traits and embraces a factor analysis model (FAM) with *K* latent factors to characterize the between-trait dependence. Additionally, since FAM in DrFARM allows implicitly for missing heritability in GWAS [27, 28], it is appealing in the analysis of pleiotropic variants. Moreover, a joint analysis of *P* variants and *Q* traits can better estimate the loading coefficients in FAM and subsequently improves both estimation and power. DrFARM also extends the sparse multivariate FARM by allowing a certain kinship structure to correlate latent factors in FAM, as opposed to independent latent factors assumed in sparse multivariate FARM. We show that FAM in DrFARM is equivalent to the specification of genetic random effects in the linear mixed model [16–20], but the former has parsimonious model constructs and thus is potentially advantageous for model interpretability.

A schematic workflow of DrFARM is given in Figure 2. To handle simultaneously many variants and traits, in Step 1, DrFARM uses the regularization technique under a sparse group lasso penalty, resulting in both individual (entry-level, i.e., all variant-trait coefficients) level and group (variant-level) level sparsity. Since the sparse estimation does not have the capacity to intentionally control any error rate (e.g. FDR) in the analysis, this method is limited for its use in GWAS when the quantification of sampling uncertainty and discovery rate control are of primary interest. Step 2 of DrFARM implements a rigorous statistical inference through the debiasing technique, leading to valid asymptotic distributions to generate desirable inferential quantities such as *p*-values and confidence intervals for individual association parameters. Step 3 of DrFARM uses the standard FDR control techniques (e.g. Benjamini-Hochberg procedure [29]) along with the Cauchy combination test (CCT) to calculate combined *p*-values for the detection of pleiotropic variants.

**Fig. 2.**
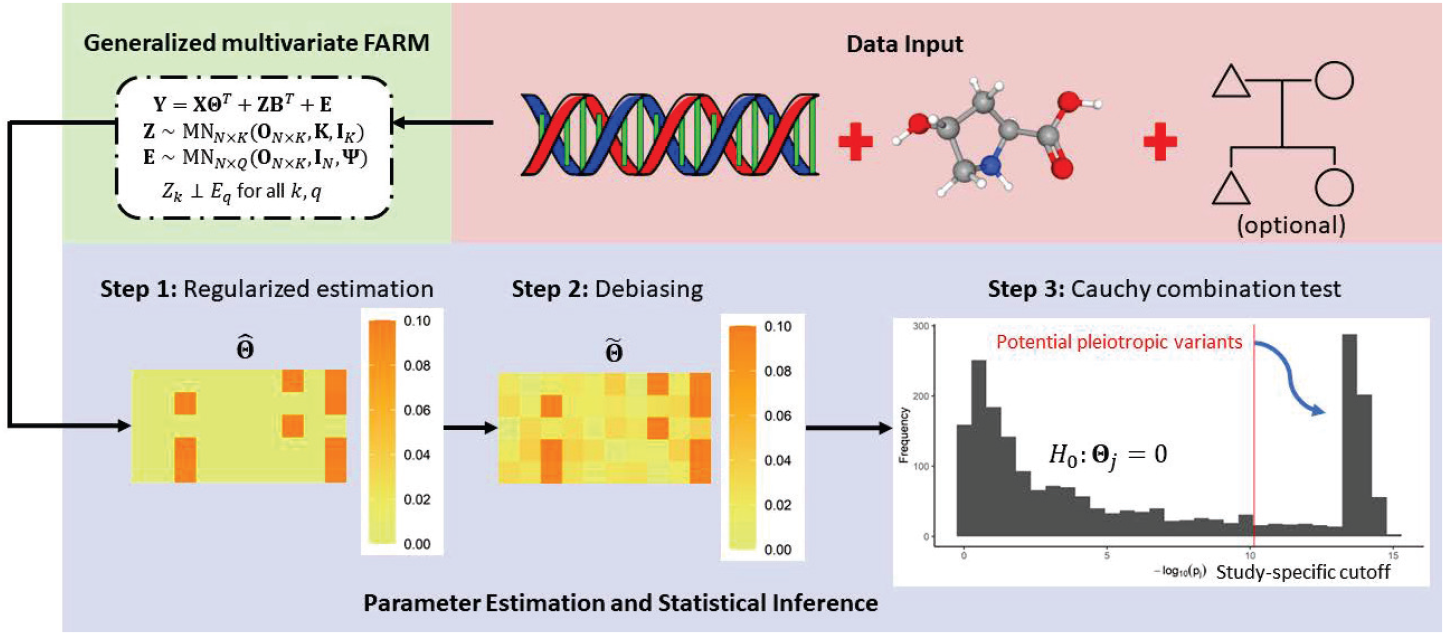
Schematic workflow of the DrFARM method with three major steps.

### 2.3 Simulation

We conduct extensive simulation experiments to evaluate the performance of the proposed DrFARM, two of which are reported in detail in this paper. The first compares the standard sparse multivariate FARM with no debiasing and three modified sparse multivariate FARM procedures with (i) only inner debiasing, (ii) only outer debiasing, and (iii) with double debiasing (i.e. both inner and outer debiasing) under various choices of precision matrix estimation methods, including Glasso, nodewise lasso, quadratic optimization and naïve (no use of the precision matrix in inner debiasing). Inner debiasing refers to a debiasing step taken in the M-step of the EM algorithm (see Algorithm 1 in Methods); outer debiasing operates a desparsifying step to ensure the asymptotic normality for individual sparse estimates. The remMap approach [15], which does not involve FAM, is also included in the comparison as the most parsimonious joint model. The second simulation investigates the influence of kinship to be or not to be included in the latent factors of FAM when data are sampled from genetically related subjects. In each simulation setting, we vary the sample size, number of SNPs, number of traits, and number of latent factors. See Table 1 for a more detailed description of simulation settings.

**Table 1.** Simulation scenarios used in experiments 1 and 2. Number of signals is defined as the number of nonzero elements in **Θ**. In each scenario, the number of pleiotropic variants (*m*) is fixed at 15% of the number of SNPs.

| Scenario | Sample Size | Predictors | Traits | Latent Factors | Signal |
| --- | --- | --- | --- | --- | --- |
| I | 1000 | 2000 | 500 | 5 | 3000 |
| II | 2000 | 5000 | 1000 | 10 | 7500 |

In simulation I, we generated data from a standard sparse multivariate FARM assuming independent individuals. As seen in Scenario I in Figure 3, all methods that do not use outer debiasing appear to have high FDRs at both individual and group-levels. Similarly, Scenario II in Table 2 suggests that both remMap and the naïve method perform poorly in the FDR control without using outer debiasing. The naïve method inflates individual-level and group-level FDRs as high as 27.2% and 65.9%, respectively.

**Fig. 3.**
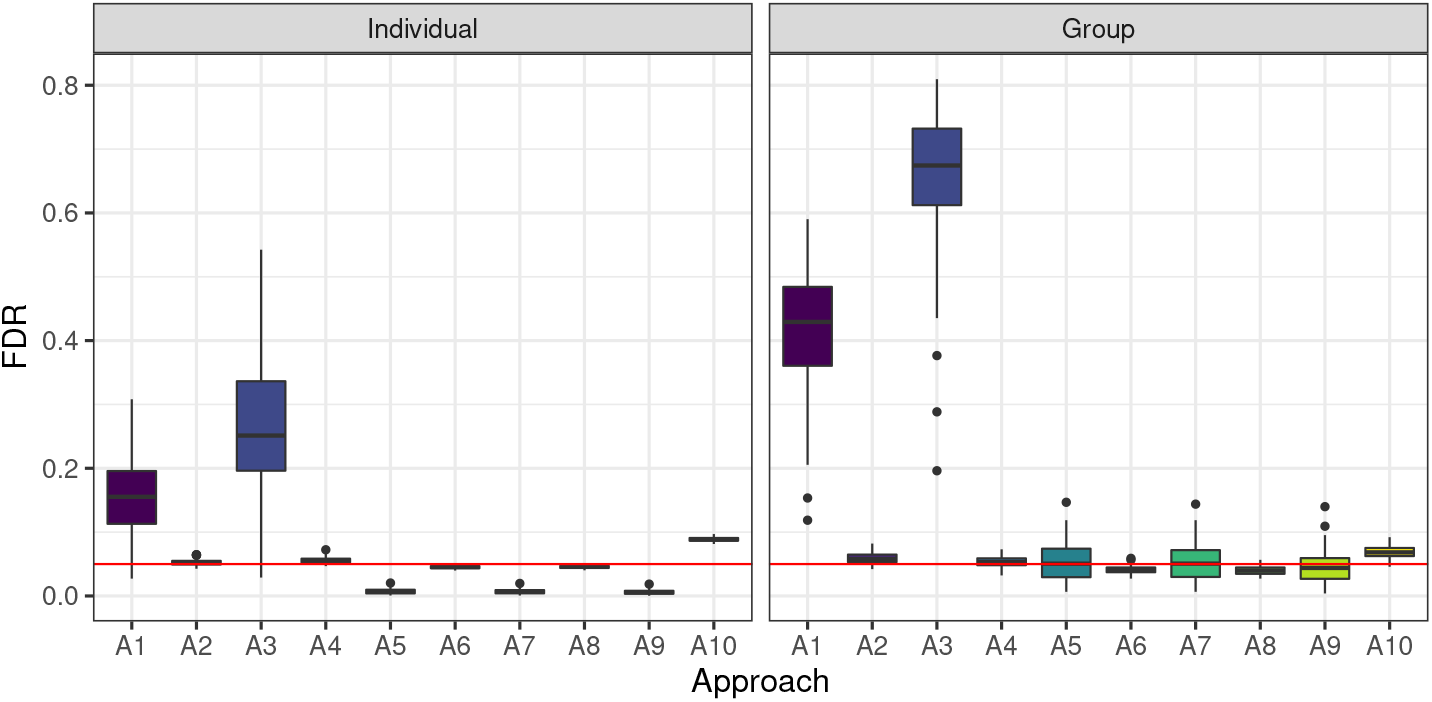
Individual-level and group-level false discovery rates for 10 different approaches (A) across 100 replicates: A1: remMap.none; A2: remMap.outer; A3: Naïve.none; A4: Naïve.outer; A5: Glasso.inner; A6: Glasso.double; A7: NL.inner; A8: NL.double; A9: QO.inner; A10: QO.double.

**Table 2.** Averaged performance metrics across 100 replicates for remMap (*r*) and DrFARM (*d*) under different type of debiasing in Scenario II for simulation 1. The true negative rate (TNR) and true positive rate (TPR) were not shown for the individual-level and group-level results, respectively, as all methods achieve close to 100%.

| Method | Precision | Debiasing | Individual |  |  | Group |  |  |
| --- | --- | --- | --- | --- | --- | --- | --- | --- |
|  |  |  | TPR | FDR | MCC | TNR | FDR | MCC |
| $d$ | None | None | 99.7% | 27.2% | 85.0% | 61.6% | 65.9% | 45.8% |
| $d$ | Glasso | Outer | 99.3% | 5.6% | 96.8% | 99.0% | 5.4% | 96.8% |
| $d$ | Glasso | Inner | 95.2% | 0.7% | 97.2% | 99.0% | 5.3% | 96.8% |
| $d$ | Glasso | Double | 98.2% | 4.6% | 96.8% | 99.2% | 4.1% | 97.5% |
| $d$ | NL | Inner | 95.2% | 0.7% | 97.2% | 99.0% | 5.3% | 96.8% |
| $d$ | NL | Double | 98.0% | 4.6% | 96.7% | 99.3% | 4.0% | 97.6% |
| $d$ | QO | Inner | 95.4% | 0.6% | 97.4% | 99.2% | 4.4% | 97.3% |
| $d$ | QO | Double | 96.4% | 8.9% | 93.7% | 98.7% | 6.8% | 95.9% |
| $r$ | None | None | 94.2% | 15.6% | 89.1% | 86.7% | 41.6% | 71.0% |
| $r$ | Glasso | Outer | 90.8% | 5.3% | 92.7% | 98.9% | 5.9% | 96.4% |

In regard to the choice of precision matrix estimation, the strategy of the inner debiasing appears to be very conservative; despite achieving accurate FDR control at 5% for the group-level signals, the FDRs for individual-level signals range from 0.6 − 0.7%. This shows that there is a conservative FDR control by the regularized method. In contrast, for the strategies involving the use of the outer debiasing, four methods (remMap, naïve, Glasso and node-wise lasso) are all able to control their FDRs at levels close to 5% for both individual-level and group-level signals, except the strategy using the quadratic optimization method the precision matrix estimation yields on average 8.9% FDR for individual signals and 6.8% FDR for group-level signals. In addition to FDR, we compare their performances by MCC (Matthews correlation coefficient), a composite metric of sensitivity and specificity. From Table 3 in Appendix D, we see that the naïve, Glasso and nodewise lasso with the outer debiasing show very similar MCCs for the detection of both individual-level and group-level signals. In Scenario I, the MCC values in Table 3 indicates that the naïve method with the outer debiasing is slightly more powerful than Glasso and nodewise lasso for the detection of both individual-level and group-level signals. In summary, outer-debiasing seems to be essential in controlling FDR while not being too conservative.

In simulation II, we simulate data by mimicking GWAS of common variants (≥5% minor allele frequency) in genetically related individuals of on average the third-degree relatedness. Based on our experiences from simulation I that no use of the outer debiasing leads to an unsatisfactory FDR control, we here only focus on the results from the methods with the utility of the outer debiasing. As shown in Figure 4. (Scenario I), the FDR for individual-level signal for the quadratic optimization method appeared constantly above 5% regardless of accounting for kinship or not whereas the FDR for group-level signals is controlled under 5%. All the other methods of precision matrix estimation exhibit satisfactory FDR control at levels close to or below 5%. In particular, the FDR for the individual-level signal was uniformly very close to 5%. Furthermore, from the performance results in terms of MCC in Tables 4 (Scenario I) and 5 (Scenario II) in Appendix D, we again observe that the naïve method, with or without kinship, is slightly more powerful than both Glasso and node-wise lasso methods for the detection of both individual-level and group-level signals. Incorporating kinship in the analysis does not lead to gains in MCC due largely to the fact that MCC is not a metric of statistical power (or one minus type II error) but a metric of detection accuracy composed by sensitivity and specificity.

**Fig. 4.**
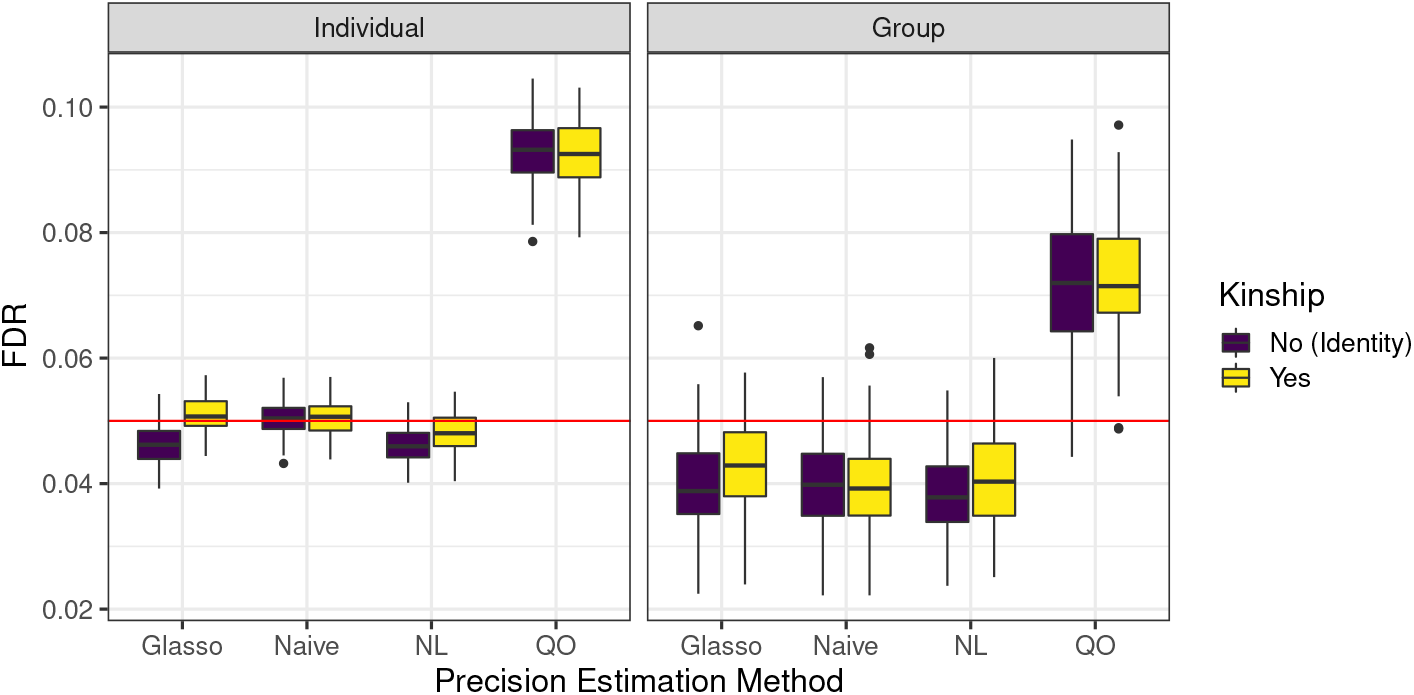
Individual-level and group-level false discovery rates obtained under 2 kinship settings by 4 precision matrix estimation approaches dealing with the outer debiasing.

In conclusion, incorporating kinship does not seem to improve FDR significantly and we recommend not using it to improve computational efficiency. In addition, among the 3 precision estimation approaches (Glasso, naïve and nodewise lasso) with FDR control, we recommend Glasso as it utilizes the inner-debiasing step and the computational complexity (or CPU time) is the lowest.

### 2.4 Real data application

Given the high correlation of metabolite abundance for many sets of metabolite across METSIM study participants, we expect to see that many loci exhibit pleiotropy across those metabolite sets. In the original single metabolite GWAS [30], we found at least one significant (*P* < 7.2 × 10^− 11^) association for 803 of the 1,031 tested metabolites. Of the 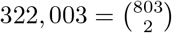 possible combinations of these metabolites, 334 have a a high phenotypic correlation (i.e., *ρ* ≥ 50%). And of the 334 highly correlated metabolite pairs, 257 (77%) exhibit pleiotropy in at least one locus, where we define pleiotropy as having significant hits for each metabolite within 10kb of each other (Supplementary Table 3, [30]). For example, the two medium chain acylcarnitines hexanoylcarnitine and octanoyl-carnitine both have significant lead SNPs at the *ACADM* locus (encoding the medium-chain acyl-CoA dehydrogenase), which was unsurprising considering this enzyme acts on both metabolites [31], and both the metabolites are strongly correlated, *ρ* = 63.6%.

Similarly, 257 (4.5%) of the 5,176 unique metabolite pairs sharing a locus (at least one significant hit for each metabolite within 10kb of each other) in [30], have a high phenotype correlation. Thus at least some of observed pleiotropy can be explained by the phenotypic correlation of the metabolite concentrations. However, a single locus can also be significantly associated with traits that are not highly correlated at the phenotypic level. For example, hexanoylglycine has a significant association at the *ACADM* locus even though the phenotypic correlation *ρ* with hexanoylcarnitine is only 18.5%.

Because DrFARM uses the correlation structure across the metabolites to enhance the power to detect genetic associations for individual metabolites, we explored the extent to which the associations identified by DrFARM reflect these phenotypic correlations. Of the 77 = 334 − 257 highly correlated metabolite pairs with no pleiotropic loci in the original study, DrFARM detected a significant association for an additional 16 of the 77. For example, the caffeine metabolites 1-methylurate and paraxanthine share a phenotypic correlation *ρ* = 57.8%, and yet while paraxanthine was significantly associated with the *CYP2A6* locus (*p* = 2.2 × 10^−19^ at rs56113850) in the single metabolite GWAS, 1-methylurate has a *p*-value of only 0.0013 at this same variant in the single metabolite analysis. In contrast, DrFARM assigns a *p*-value of 3.9 × 10^−13^ to 1-methylurate at rs56113850. This association is highly plausible given that the CYP2A6 enzyme is responsible for acting on paraxanthine on its way to being converted to 1-methylurate.

In all, DrFARM assigned a *p*-value < 7.2 × 10^−11^ to 288 metabolite-locus pairs where the prior metabolite GWAS analysis had no significant association for that specific metabolite within 100,000 bps. While the new metabolite associations are skewed toward metabolites that are highly correlated with the previously identified metabolites, 70% of the new metabolite associations does not have high correlation to any of the previous metabolites at the locus. For example, at the *GLS2* locus (encoding a glutaminase enzyme) the single metabolite GWAS identified significant associations for both glutamine and a glutamine derivative, gamma-glutamylglutamine. DrFARM found an additional association for another glutamine derivative, hexanoylglutamine, despite the fact that hexanoylglutamine and glutamine share a phenotypic correlation (*ρ*) of only 0.06%. Despite the low phenotypic correlation of most of the new metabolite associations from DrFARM compared to the previous single metabolite results, the vast majority of the new results represent highly plausible biological results. For example, where the previous analysis identified tyrosine as a significant association at the *TAT* locus (encoding tyrosine aminotransferase), the new analysis identified a significant association for the tyrosine derivative, N-acetyltyrosine. The new analysis also identified a significant association for kynurenine at the *KMO* locus (encoding kynurenine 3-monooxygenase), for the caffeine derivatives 1-methylurate, 3,7-dimethylurate, 1,7-dimethylurate at the *CYP2A*6 locus (encoding a caffeine metabolizing enzyme), for the pyrimidine metabolite uracil at the *CDA* locus (encoding the pyrimidine metabolizing enzyme, cytidine deaminase) and the very long acyl carnitine 5-dodecenoylcarnitine at the *ACADVL* locus (encoding the very long-chain specific acyl-CoA dehydrogenase). Cross-referencing the DrFARM detected significant associations with biological knowledge gleaned from the rich history of biochemistry provides independent validation of these results. Expanding the current analysis to identifying pleiotropic genes for multiple metabolites is a future research direction.

## 3 Discussion

We developed a new method, DrFARM, to identify potential pleiotropic variants in GWAS. Our methodological contribution centers on one-stage post selection hypothesis testing, adjusting for other genetic variants and confounding factors. DrFARM provides satisfactory FDR control in the detection of both individual-level (entry-level) and group-level (variant-level) signals. In addition, DrFARM incorporates population structure in the latent factors as part of the modeling of between-trait correlations. Being a nontrivial extension from low-dimensional joint modeling approach, DrFARM overcomes a difficult problem of proper FDR control in the large-*P* -small-*N* setting, which has troubled existing pairwise single-variant marginal association testing in the GWAS literature.

DrFARM provides a principled approach to perform a refined downstream analysis, such as colocalization. Even though we used the set of index variants as the input genetic markers for the METSIM data analysis, following the identification of potential pleiotropic variants, we could further identify the corresponding putative causal gene of the variant and construct a respective gene region corresponding to the putative causal gene. Instead of constructing gene regions from potentially spurious variants (due to LD), DrFARM enables us to identify a more reliable and promising candidate gene regions for downstream analysis using potential pleiotropic variants.

A proven advantage of DrFARM is that it can increase power by taking into account the correlation between related traits, enabling identification of association not identified in single trait analyses. We identified 16 new candidate genes with DrFARM in the METSIM data analysis. DrFARM is not limited to the association study of metabolites-genetic variants but is applicable to other high-dimensional omics data types such as proteins and glycans. Thus, DrFARM presents an ample opportunity to discover pleiotropic variants in the integrative analysis of multi-trait and multimodal omics data in the modern biology era.

DrFARM has some limitations that deserve further exploration in future research. First, DrFARM is built upon *L*_1_ penalty regularization which is known to suffer from overfitting when predictors are highly correlated. We have seen the sensitivity of FDR on modest or highly correlated SNPs (e.g., correlation ≥ 0.7), indicating a need to invoke a better regularization method to improve DrFARM with correlated SNPs. Second, DrFARM requires the use of an estimated precision matrix in the outer debiasing step to calculate *p*-values for inference. Taking our recommended method Glasso (balancing computational efficiency and statistical performance) as an example, the computational complexity is *O*(*P* ^3^) to *O*(*P* ^4^), depending on the actual sparsity of the precision matrix [32]. Thus, DrFARM is computationally expensive to handle tens of thousands of variants, which might be improved by feature screening methods [33] to reduce dimensionality prior to the application of DrFARM, or by a fast precision matrix estimation method.

As for future work, one direction is to investigate the latent factors used by DrFARM. Similar to traditional factor analysis, the interpretation of latent factors is a challenging issue. Potentially, geneticists could mine the latent factors to understand the missing heritability in GWAS, similar to how principal component analysis (PCA) has helped to understand population stratification [34]. Related tasks would include associating these latent factors with different gene regions and elucidating what kind of factor rotation provides a meaningful interpretation for the latent factors. With the ever-increasing size of GWAS cohorts and whole genome sequencing platforms, another important work is to develop scalable algorithms for estimating ultra high-dimensional precision matrices as they play a crucial role in statistical inference with high-dimensional genomics data.

## 4 Tables

## 5 Figures

## Methods

### 1 Setup in motivating example

Consider two correlated traits, *Y*_1_ and *Y*_2_, constituting a bivariate trait by **Y** = (*Y*_1_, *Y*_2_). Suppose that **Y** is generated from the true model

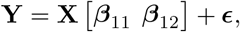

where **X** = (*X*_1_, · · ·, *X*_*P*_) is a set of *P* predictors (e.g., SNPs), ***β***_11_ and ***β***_12_ are *P* -dimensional vector of true coefficients associating **X** with *Y*_1_ and *Y*_2_, respectively (notice that some of the coefficients of ***β***_11_ and ***β***_12_ can be zero). Since the traits are correlated, we assume a phenotypic correlation *ρ*, for Var(***ϵ***), where *ρ* ≠ 0.

In practice, it is often assumed that the *P* SNPs are independent and contribute to the traits independently. However, this assumption may be violated for genetics data due to factors including linkage disequilibrium and population structure [35].

We set *N* = 6135, *P* = 2072 (same as our real data analysis setting) and suppose there are 250 true SNPs that contribute to the two traits. The effect sizes of true SNPs are generated by sampling 500 = 250 × 2 effect sizes from the set of 3443 genomewide significant associations from prior METSIM single metabolite GWAS. [30]. We also set a weak phenotypic correlation *ρ* = 0.3. SNPs are generated by sampling 2072 SNPs from a set of 6334 LD-pruned SNPs from chromosome 22 using METSIM data with *r*^2^ = 0.01 threshold. The empirical type I error is given by the number of significant discoveries (i.e., p-value < 0.05) in the null set divided by 1822 = 2072 − 250 (the number of null), which is evaluated from 1000 replicates.

### 2 Review of remMap and sparse multivariate FARM

Both remMap and sparse multivariate FARM are regularized multivariate regression models that exploit sparse group lasso penalty to identify “master” predictors (i.e., pleiotropic variants in GWAS). In particular, sparse multivariate FARM extends remMap by modeling residual correlations of traits via a latent factor model [13]. More specifically, assume *P* SNPs and *Q* traits are collected in each individual. Let **x**_*i*_ = (*x*_*i*1_, *· · ·, x*_*iP*_)^*T*^ and **y**_*i*_ = (*y*_*i*1_ *· · ·*,, *y*_*iQ*_)^*T*^ (*i* = 1, …, *N*) be normalized SNPs and normalized traits with mean 0 and variance 1, respectively. The multivariate FARM takes the form:

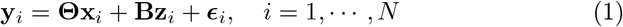

where **Θ** = *θ*_*qp*_ is a *Q* × *P* coefficient matrix, **B** is a *Q* × *K* matrix of factor loadings (*K* being the number of latent factor). Multivariate FARM assumes the latent factors **z**_*i*_ = (*z*_*i*1_, · · ·, *z*_*iK*_)^*T*^ ∼ MVN_*K*_(**0**_*K*_, **I**_*K*_). Moreover, ***ϵ***_*i*_ = (*ϵ*_*i*1_, · · ·, *ϵ*_*iQ*_)^*T*^ ‘s are independent and identically distributed (i.i.d.) errors from MVN_*Q*_(**0**_*Q*_, **Ψ**) with **0**_*Q*_ being a *Q*-element zero vector and **Ψ** = diag(*ψ*_1_,, *ψ*_*Q*_) being a *Q* × *Q* diagonal matrix. The multivariate FARM further assume ***ϵ***_*i*_ is independent of the latent factors **z**_*i*_.

The multivariate FARM has the following equivalent form:

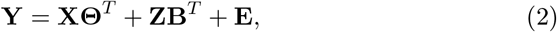

where **Y**_*N* × *Q*_ = (**y**_1_, · · ·, **y**_*N*_)^*T*^, **X**_*N* × *P*_ = (**x**_1_, · · ·, **x**_*N*_)^*T*^, **Z**_*N* × *K*_ = (**z**_1_, · · ·, **z**_*N*_)^*T*^ ∼ MN_*N* × *K*_(**O**_*N* × *K*_, **I**_*N*_, **I**_*K*_) and **E**_*N* × *Q*_ = (***ϵ***_1_, · · ·, ***ϵ***_*N*_)^*T*^ ∼ MN_*N* × *Q*_(**O**_*N* × *Q*_, **I**_*N*_, **Ψ**). Here MN_*n* × *m*_(**M, V**_*r*_, **V**_*c*_) denotes the *n* × *m* **m**atrix **n**ormal distribution with mean matrix **M** (*n* × *m*), row (inter-sample) covariance matrix **V**_*r*_ (*n* × *n*) and column (between component) covariance **V**_*c*_ (*m* × *m*). The conditional covariance of the response variables given the predictors is Var(**y**_*i*_|**x**_*i*_) = **Σ** = **BB**^*T*^ + **Ψ**.

The objective function of sparse multivariate FARM is given by

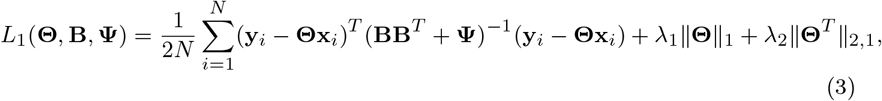

where 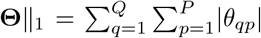 and 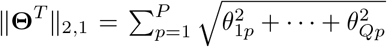, and *λ*_1_, *λ*_2_ > 0 are tuning parameters controlling the entrywise sparsity and column-wise sparsity in **Θ**, respectively.

We estimate the parameters (**Θ, B, Ψ**) in sparse multivariate FARM using the EM-GCD algorithm [13], which uses a group-wise coordinate descent (GCD) algorithm for estimating **Θ** and expectation-maximization (EM) algorithm for estimating both **B** and **Ψ**. When there are no latent factors (i.e., *K* = 0), Model (1) reduces to the remMap model. The objective function of remMap is given by

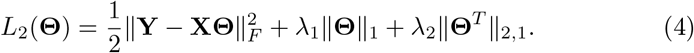

Notice that (4) implicitly assumes the variance of the *Q* trait residuals are equal. The parameter **Θ** is estimated using a modified version of the active shooting algorithm [15, 36, 37]. More details of remMap and sparse multivariate FARM may be found in [15] and [13], respectively.

### 3 Generalized multivariate FARM

We consider a generalization of the multivariate FARM in DrFARM where the latent factors are allowed to be correlated when study participants are related. That is, we specify **Z** ∼ MN_*N* × *K*_(**O**_*N×K*_, **K, I**_*K*_), where **K** (*N* × *N*) is a prespecified kinship matrix that is scaled to have diagonal 1 analogous to a correlation matrix. In GWAS, **K** is typically estimated separately from available genotype data, e.g., using KING [38]. To decorrelate samples, we perform an eigendecomposition of **K** = **UDU**^*T*^ [17, 20, 39, 40], where **U** is an *N* × *N* orthogonal matrix of eigenvectors and **D** = diag(*δ*_1_, *· · ·, δ*_*N*_) is an *N* × *N* diagonal matrix of eigenvalues. Correspondingly, an equivalent form of the generalized multivariate FARM is

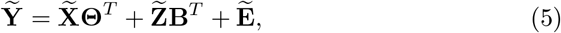

where 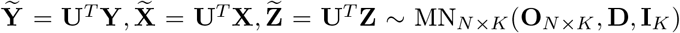 and 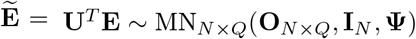. That is, for each individual *i*,

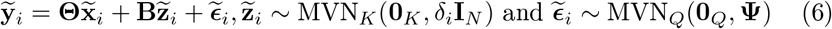

where 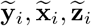 and 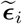 are the *i*th row of 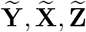 and 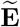 respectively. Note that there is an extra *δ*_*i*_ term in the variance of 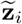 compared to **z**_*i*_ in (1) due to the presence of kinship dependence among subjects. With the transformation, the likelihood can be obtained as a product of *N* individual likelihoods, which can be easily evaluated. To deal with latency of 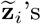, we invoke the EM algorithm by treating the 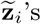 as missing data in the estimation of the model parameters (**Θ, B**).

The generalized multivariate FARM connects to the multivariate linear mixed model GEMMA given in [40]:

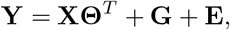

where **G**_*N* × *Q*_ ∼ MN_*N* × *Q*_(**O**_*N* × *Q*_, **K, V**_*g*_) is genetic random effects, **E** ∼ MN_*N* × *Q*_(**O**_*N* × *Q*_, **I**_*N*_, **V**_*e*_), **V**_*g*_ is the *Q* × *Q* symmetric matrix of genetic variance component and **V**_*e*_ is the *Q* × *Q* symmetric matrix of environmental variance components. In comparison, generalized multivariate FARM is more parsimonious by modeling the random effects **G** with FAM **ZB**^*T*^ ∼ MN_*N* × *Q*_(**O**_*N* × *Q*_, **K, BB**^*T*^) (or equivalently, **V**_*g*_ = **BB**^*T*^). FAM presents simpler covariance structures to both genetic and environmental variance component matrices, and the latent factors may be used to investigate the missing heritability in GWAS (see Discussion).

### 4 Regularized estimation

The complete data log-likelihood is

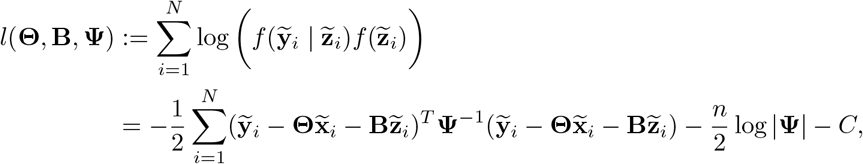

where *C* is a constant.

To identify pleiotropic variants, we employ a regularized estimation method via the sparse group lasso penalty (by predictor/column) *λ*_1_ ∥**Θ**∥ _1_ + *λ*_2_∥ **Θ**^*T*^∥_2,1_ to achieve sparse estimation of **Θ**, where *λ*_1_, *λ*_2_ are tuning parameters controlling the entrywise sparsity and column-wise sparsity in **Θ**, respectively. This penalized estimation is integrated with the EM algorithm that deals with the augmented data log-likelihood with latent factors **Z**. The penalized log-likelihood function for complete data is given by

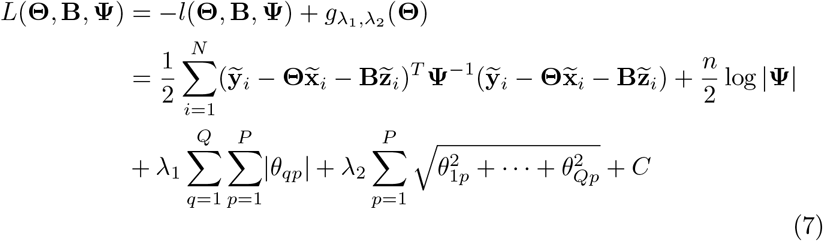

where *g*_*λ, λ*_ (**Θ**) := *λ*_1_∥**Θ**∥_1_ + *λ*_2_∥**Θ**^*T*^ ∥_2,1_ and *C* is a suitable constant with respect to the parameters (**Θ, B, Ψ**).

Let *t* be the iteration number. In the E-step we calculate the first two conditional moments

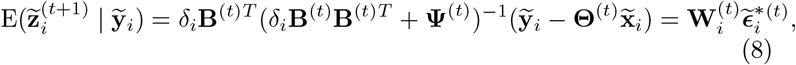

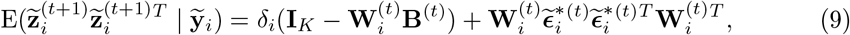

where **W**_*i*_ = *δ*_*i*_**B**^*T*^ (*δ*_*i*_**BB**^*T*^ + **Ψ**)^−1^ and 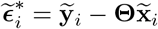.

In the M-step, we compute 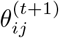 (see expression (15)),

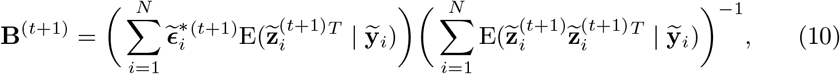

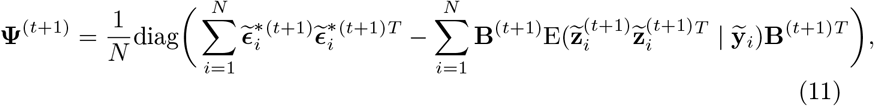

For the detailed derivation, please refer to Appendix A. Let 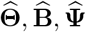 be the regularized estimator for **Θ**, EM estimator for **B** and **Ψ**, respectively. Also, let 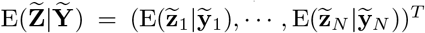. Then, we denote the conditional moment based on estimators 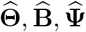 by 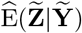. Define 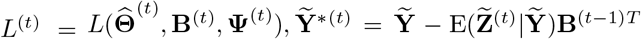 and 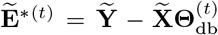. The pseudocode of the EM algorithm for parameter estimation is given in Algorithm 1. We highlight two major differences compared to the algorithm implemented in sparse multivariate FARM [13]: (i) Instead of obtaining an exact minimizer of 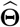 in **M-step 1**, we use a one-step update [41] to reduce the computational cost. Our numerical studies show that the one-step approximation does not change the final estimate much but greatly improves the overall computational efficiency. (ii) We add a second **M-step 2** to calculate a debiased estimate 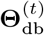. This debiasing step helps us to get a more stable estimate of the residual matrix 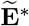, which subsequently enhances the estimation of the quantities in the FAM (**B, Ψ**) in **M-step 3**. We refer to **M-step 2** as **inner debiasing**. The initial value determination and tuning parameter selection are detailed in the Appendix C.

### 5 Estimation of variance parameters

The estimates of the trait residual variance (or uniqueness) *ψ*_*i*_ (for *i* = 1, …, *Q*) are part of the parameters output from the EM algorithm. The true *ψ*_*i*_’s are typically underestimated in numerical studies. As a remedy, we propose an alternative estimator adjusting for the degrees of freedom given by

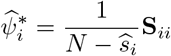

where

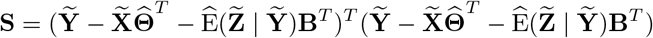

and 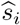 is the number of nonzero in the *i*th row of 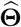 (i.e., all the coefficients associated with trait *i*). Likewise, estimator of variance *σ*^2^ is given by

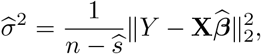

which is suggested by [42] (Section 2.2), *s* is the number of nonzero in the lasso estimator 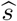.

### 6 Inference

#### 6.1 Single parameter inference

In the univariate regression analysis *Y* = **X*β*** + *ϵ* (*ϵ ∼ N* (0, *σ*^2^)), a lasso estimator 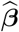 [43] can be desparsified (termed in [21]) or debiased (termed in [23]) by

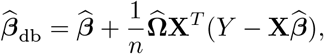

where

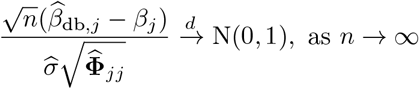

under some regularity conditions, 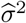 is an estimator for σ ^2^ when *n* < *p* (see Section 5). In particular, 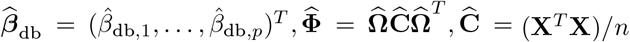, and 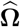 is the estimated precision matrix which approximates *n*(**X**^*T*^ **X**)^−1^ when *n < p*.

In the same spirit, we propose to debias the regularized estimator 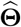 in DrFARM by

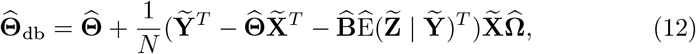

where 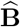 and 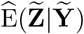are estimators of **B** and 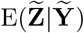 obtained from the EM algorithm (see Appendix A). Correspondingly, similar asymptotic properties can be derived for 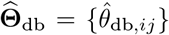 (see Appendix B). We refer to this as an **outer debiasing** step. The outer-debiasing step is different from the **inner-debiasing** step, which is used **inside** the EM algorithm. The outer-debiasing step is used **outside** of the EM algorithm (once the estimation is completed) for statistical inference. Despite the difference in purpose, the outer and inner debiasing steps share a common debiasing expression. It follows that the *p*-value for testing *H*_0_ : *θ*_*ij*_ = 0 involving the *i*th trait and *j*th predictor *p*_*ij*_ can be calculated by the above estimator with

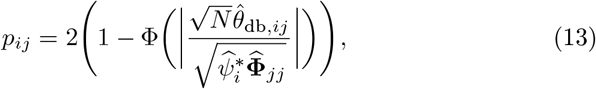

where 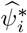 is an estimator for uniqueness (see Section 5) and Φ is the cdf of the standard normal distribution.

#### 6.2 Hypothesis test for pleiotropy

Let **Θ**_*j*_ be the *j*th column of **Θ**. Testing for pleiotropy (also known as testing the **group-level** significant association) is equivalent to testing **Θ**_*j*_ = 0. Of note, the classical MANOVA test statistics, such as Wilk’s Lambda [44], Pillai’s Trace [45], Hoteling-Lawley Trace [46] and Roy’s Greatest Root [47] cannot be used when *P > N*. To use the asymptotic result in [48], we consider the Cauchy combination test (CCT) [48] for the joint test of **Θ**_*j*_ = 0. The CCT takes the form

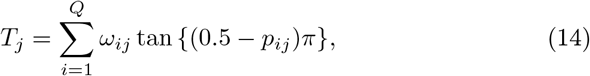

where *ω*_*ij*_ are nonnegative weights and 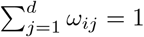. The test statistic follows a Cauchy distribution under the null with an arbitrary dependence structure between *p*_*ij*_’s. Liu and Xie demonstrated that CCT can be used for single trait discovery in GWAS [48]. For our purpose, we extend the CCT to multitrait discovery and adjust for multiple testings using the Benjamini-Hochberg procedure [29]. More specifically, we obtain individual *p*-value *p*_*ij*_ using (14) and plug it in the CCT test statistic formula. The corresponding *p*-value *p*_*j*_ is then given by

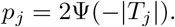

where Ψ is the cdf of the standard Cauchy distribution.

### 7 Choice of precision matrix estimation

The precision matrix plays a critical role in the debiasing steps. There is a large body of literature on precision matrix estimation. However, to the best of our knowledge, the influence of different estimation methods on the statistical performance of the debiased estimator [21–23] has not been studied. Here we compare three precision matrix estimation methods: 1) Graphical Lasso (Glasso) maximizes the penalized log-likelihood [26] but with unknown theoretical guarantees [21]; 2) Nodewise lasso (NL), performs row-wise lasso and proved theoretical guarantees in estimation consistency [21] and 3) Quadratic optimization (QO) performs a row-wise convex optimization with theoretical guarantees in estimation consistency [23].

In our numerical studies, we exploited the precision matrix estimated from Glasso and NL where tuning parameters were selected by the extended Bayesian information criterion (EBIC) with *γ* = 0.5 [49, 50]. For Glasso, we used 10 tuning parameters (default setting) using `glassopath()` of the R package `glasso`. In the same spirit, for NL, we fitted *P* regression models *X*_*i*_ regressed on **X**_−*i*_ for all *i* = 1, …, *P* (where *X*_*i*_ denotes the *i*th column of **X** and **X**_−*i*_ denotes the matrix after omitting *i*th column from **X**) and used 100 tuning parameters (default setting) using R package `glmnet`. For QO, we used the R code provided on the first authors’ website: https://web.stanford.edu/~montanar/sslasso/code.html with the default setting.

## 8 Simulation

In each setting, sample size (*N*), number of predictors (*P*), number of traits (*Q*), number of latent factors (*K*), and number of signals are all varied. We implement the proposed method and use EBIC (*γ* = 1) for tuning parameter selection. We use 100 replicates for all the methods compared. Details for the implementation of the methods can be found in Appendix C.

### 8.1 Simulation I

Suppose **X** = {*x*_*np*_}, **Z** = {*z*_*nk*_} and **E** = {*ϵ*_*nq*_}. Their entries *x*_*np*_, *z*_*nk*_ and *ϵ*_*nq*_ are independently generated from N(0, 1) for *n* = 1, …, *N, p* = 1, …, *P, k* = 1, …, *K* and *q* = 1, …, *Q*. To generate the *Q×P* coefficient matrix **Θ** = {*θ*_*qp*_} between the *Q* traits and *P* predictors, we specify a sparse indicator matrix **Δ** = *{δ*_*qp*_*}*. If *δ*_*qp*_ = 1, then *θ*_*qp*_ ∼ Unif([−1.5, −1]∪[1, 1.5]). Otherwise, *θ*_*qp*_ = 0. Notice that 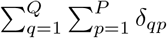 is the number of signals fixed in a given scenario. Given a fixed number of pleiotropic variant *m* (set to be 15% of the number of predictors), the set of pleiotropic variants is randomly drawn from the indices {1, …, *P*} without replacement. Let *M* = *q* : *θ*_*pq*_ = 1, for *q* = 1, …, *Q*, i.e., the set of indices corresponding to the pleiotropic variants. The number of trait associated with each *j ∈ M* follows Multinomial 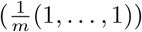. To specify the factor loading matrix **B**, we adopt an approach similar to [13]. First, we start with an initial matrix 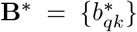 where 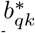 are independently generated from Unif(0, *τ*) where *τ >* 0 is determined empirically and fulfills the signal-to-signal-to-noise ratio (SSNR) = mean(diag(Cov(**XΘ**^*T*^))) : mean(diag(Cov(**ZB**^*T*^))) : mean(diag(Cov(**E**))) = 1 : 3 : 5. This SSNR is used to mimic the missing heritability scenario of GWAS and gives the necessity for modeling the latent factors. We perform an eigendecomposition **B**^*^**B**^\**T*^ = **U**^*^**Σ**^*^**U**^\**T*^ where the column vectors of **U**^*^ are orthonormal eigenvectors of **B**^*^**B**^\**T*^ and **Σ**^*^ is a diagonal matrix with diagonal entries being the eigenvalues of **B**^*^**B**^\**T*^. Then we can let **V**^*^ = sqrt(**Σ**) and form **B** = **U**^*^**V**^*^. Finally, the data are generated using the equation **Y** = **XΘ**^*T*^ + **ZB**^*T*^ + **E**.

### 8.2 Simulation II

For this simulation, all settings are kept the same as Simulation I except *x*_*ni*_ ∼ Bin(2, *p*_*i*_) independently for all *n* = 1, …, *N* and *Z*_*k*_ ∼ MVN_*N*_ (**0, K**) independently for *k* = 1, …, *K*, where **Z** = [*Z*_1_, …, *Z*_*K*_]. To mimic common variants in GWAS, *p*_*i*_ ∼ Unif(0.05, 0.95) independently for all *i* = 1, …, *P*. We generated kinship **K** using the standardized **X**^*^**X**^\**T*^ (i.e., `cov2cor()` in R) where 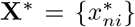 has its entries 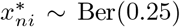∼ Ber(0.25) for *n* = 1, …, *N* and *i* = {1, …, *P*} so that the off-diagonal entries of **K** has a mean of 0.25 to simulate a third-degree relationship (2 × 0.125) between individuals on average [38].

## 9 Performance metrics

We used true positive rate (TPR), true negative rate (TNR), false discover rate (FDR) and Matthew’s correlation coefficient (MCC) [51]

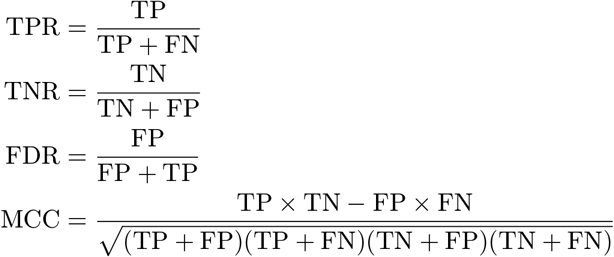

to compare the performance of different approaches in simulations I and II, at both the individual level and group (SNP) level. In particular, for methods that do not provide *p*-values (i.e., without debiasing or with inner debiasing only), the number of true positive (TP) is the number of nonzero elements in the selected 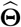 in the signal set for signal-level result and the number of pleiotropic variants with at least one nonzero association for the group (SNP) level result. The number of true negatives (TN) is the number of zeros in the selected 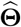 in the non-signal set for signal-level result and the number of the non-pleiotropic variant with no association for the group-level result. Then, the number of false positives (FP) and the number of false negatives (FN) simply given by the number of positive (nonzero coefficients) minus TP, and the number of negatives (zero coefficients) minus TN, respectively. For methods that provide *p*-values (i.e., outer debiasing or double debiasing), we applied Benjamini Hochberg procedure [29] to both the signal-level and group-level *p*-values at 5% level. To calculate TP, TN, FP and FN, instead of evaluating whether the coefficients are nonzero, we consider whether the adjusted *p*-values is smaller than 0.05.

## 10 METSIM dataset

We use the same metabolomics GWAS data set as in [30] to demonstrate the performance of the proposed methods. The sample size is *N* = 6135. We focused on a subset of *P* = 2072 nearly-independent index variants identified from univariate analysis after Bonferroni correction (*P* < 7.2 × 10^−11^) [30]. We chose the set of index variants because they were the most likely candidate for pleiotropic variants. As shown in [30], 27.2% of the index variants were associated with more than 2 metabolites using a single-variants association testing approach. Since multivariate regression requires a complete data matrix for traits, we focused on *Q* = 1031 targeted metabolites that were either complete or imputable using the K-nearest neighbors approach (with 5 neighbors). Examples of non-imputable metabolites include those that were only present ≤ 3 out of 4 Metabolon panels (data collected at different times). As in [30], we regressed the Metabolon-reported metabolite level on covariates (age at sampling, Metabolon batch, and lipid-lowering medication use status for lipid traits only). To obtain covariate-adjusted metabolites with mean 0 and variance 1, we inverse normalized the residuals from the regression model [30]. We based the K-nearest neighbor imputation on the inverse-normalized scale. For further details, such as data preprocessing, please refer to [30].

## 11 METSIM data analysis

We first searched a 10 × 10 tuning parameter grid and picked the optimal tuning parameters using EBIC (*γ* = 1) for remMap. Then, remMap estimates with the selected tuning parameters were used as the initial value for DrFARM to find the optimal tuning parameters from a refined 5 × 5 grid. As suggested by the simulation, we used DrFARM with double debiasing with Glasso for discovery. We varied *K* = 1 to 100 (i.e., 5 × 5 × 100 = 2500 grids were searched in total). For a fixed *k ∈ {*1, …, 100*}*, the tuning parameter was selected among the 5 × 5 grid. Since we observed EBIC decreases almost monotonically with *k*, to avoid overfitting, the residual matrix 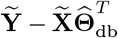 were calculated for each *k* for the selected tuning parameter. The exploratory graph analysis (EGA) [52] uses Glasso [26] to obtain the sparse inverse covariance matrix for the outcomes of interest and identifies the number of clusters or communities in a graph using a walktrap algorithm [53]. The number of dense subgraphs (communities or clusters) is declared as the number of latent factors *K*. Since metabolites are known to be clustered, we used EGA as opposed to common latent factors determination methods such as parallel analysis [54, 55] or Kaiser-Guttman’s eigenvalue-greater-than-one rule [56] for biological interpretability. We performed EGA for each of the 100 residual matrices, and majority voting of the EGA results yielded *K* = 16. The signal and SNP (group) level results were subjected to *P* < 7.2 × 10^−11^ (same cutoff as the original study) for statistical significance. Unlike simulation, in addition to *P* < 7.2 × 10^−11^ at the group-level, we also required the significant SNP to have at least 2 associated metabolites with *P* < 7.2 × 10^−11^ to be considered a **potential** pleiotropic variant.

### 12 Algorithm

EM Algorithm for a given pair of tuning parameters (*λ*_1_, *λ*_2_)

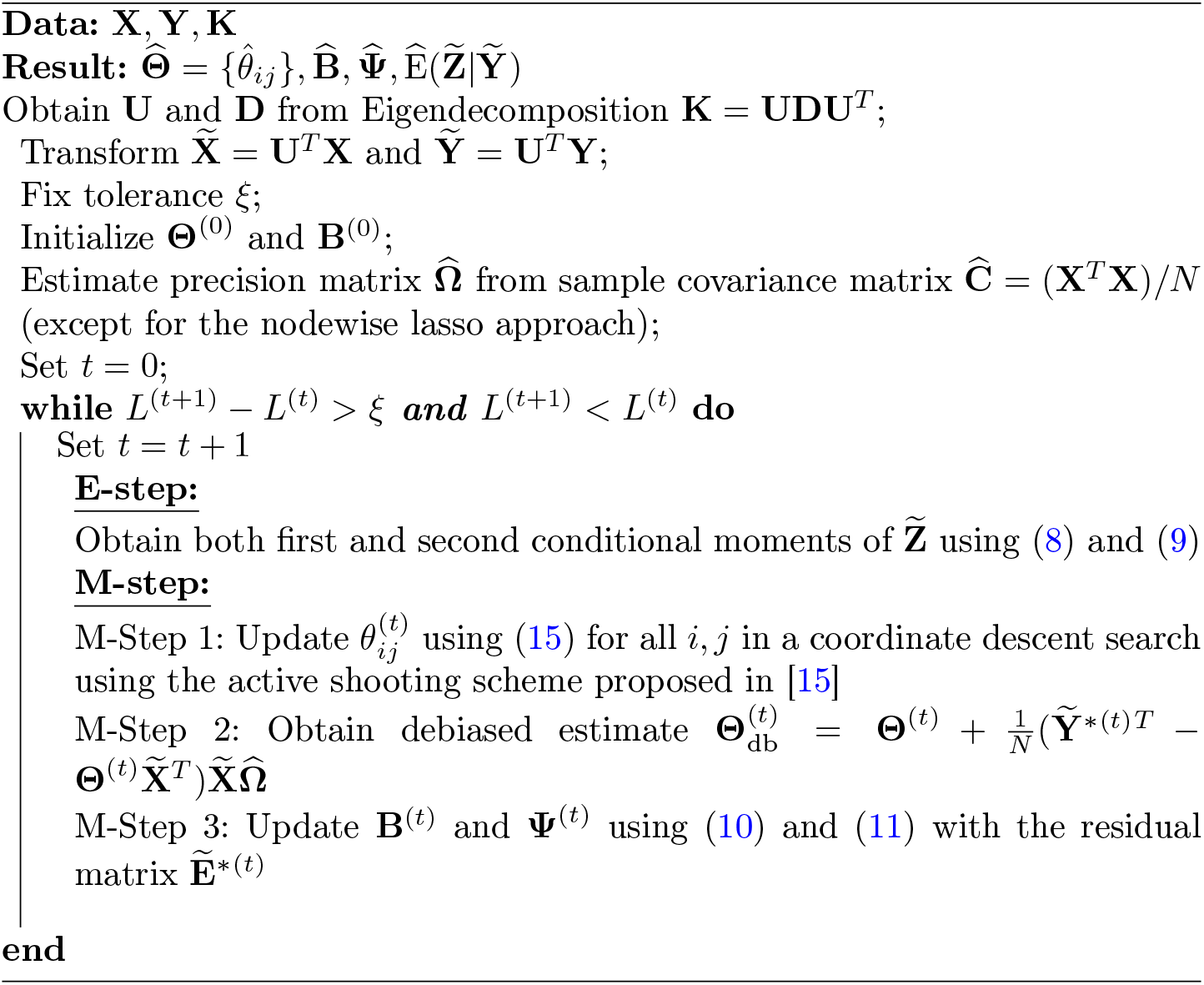

