## Appendix for "DrFARM: Identification and inference for pleiotropic gene in GWAS"

### Part III
### Appendix

#### A Derivation of EM algorithm

We consider the following model, a generalized mFARM:

$$\widetilde{\mathbf{y}}_i = \boldsymbol{\Theta}\widetilde{\mathbf{x}}_i + \mathbf{B}\widetilde{\mathbf{z}}_i + \widetilde{\boldsymbol{\epsilon}}_i, \quad i = 1, \cdots, N$$

where $\widetilde{\mathbf{y}}_i \sim \text{MVN}_Q(\boldsymbol{\Theta}\widetilde{\mathbf{x}}_i, \delta_i\mathbf{B}\mathbf{B}^T + \boldsymbol{\Psi})$.
Treating $\mathbf{z}_i$'s as missing data and we employ the EM algorithm to obtain the penalized estimate $\widehat{\boldsymbol{\Theta}}$.
Noting that $\widetilde{\mathbf{z}}_i \sim \text{MVN}_K(\mathbf{0}, \delta_i\mathbf{I})$. We have joint distribution of the form:

$$\begin{bmatrix}\widetilde{\mathbf{y}}_i \\ \widetilde{\mathbf{z}}_i\end{bmatrix} \sim \text{MVN}_{Q+K}\left( \begin{bmatrix}\boldsymbol{\Theta}\widetilde{\mathbf{x}}_i \\ \mathbf{0}\end{bmatrix}, \begin{bmatrix}\delta_i\mathbf{B}\mathbf{B}^T + \boldsymbol{\Psi} & \delta_i\mathbf{B} \\ \delta_i\mathbf{B}^T & \delta_i\mathbf{I}_K\end{bmatrix} \right)$$

due to the fact that $\text{Cov}(\widetilde{\mathbf{y}}_i, \widetilde{\mathbf{z}}_i) = \text{Cov}(\boldsymbol{\Theta}\widetilde{\mathbf{x}}_i + \mathbf{B}\widetilde{\mathbf{z}}_i + \widetilde{\boldsymbol{\epsilon}}_i, \widetilde{\mathbf{z}}_i) = \text{Cov}(\mathbf{B}\widetilde{\mathbf{z}}_i, \widetilde{\mathbf{z}}_i) = \mathbf{B}\text{Var}(\widetilde{\mathbf{z}}_i) = \delta_i\mathbf{B}$. Recall that the penalized log-likelihood function for complete data is given by

$$\begin{aligned} L(\boldsymbol{\Theta}, \mathbf{B}, \boldsymbol{\Psi}) &= -l(\boldsymbol{\Theta}, \mathbf{B}, \boldsymbol{\Psi}) + g_{\lambda_1,\lambda_2}(\boldsymbol{\Theta}) \\ &\propto \frac{1}{2}\sum_{i=1}^{N}(\widetilde{\mathbf{y}}_i - \boldsymbol{\Theta}\widetilde{\mathbf{x}}_i - \mathbf{B}\widetilde{\mathbf{z}}_i)^T\boldsymbol{\Psi}^{-1}(\widetilde{\mathbf{y}}_i - \boldsymbol{\Theta}\widetilde{\mathbf{x}}_i - \mathbf{B}\widetilde{\mathbf{z}}_i) + \frac{n}{2}\log|\boldsymbol{\Psi}| \\ &+ \lambda_1\sum_{q=1}^{Q}\sum_{p=1}^{P}|\theta_{qp}| + \lambda_2\sum_{p=1}^{P}\sqrt{\theta_{1p}^2 + \cdots + \theta_{Qp}^2} \end{aligned}$$

where the equality of $a$ to $b$ subject to a constant may be written as $a \propto b$.
In the E-step, we take the expectation of $L(\boldsymbol{\Theta}, \mathbf{B}, \boldsymbol{\Psi})$ with respect to the conditional distribution of $\widetilde{\mathbf{z}}_i|\widetilde{\mathbf{y}}_i$. Within the multivariate normality, it is sufficient to derive the two moment statistics. The conditional mean is

$$\text{E}(\widetilde{\mathbf{z}}_i \mid \widetilde{\mathbf{y}}_i) = \mathbf{0} + (\delta_i\mathbf{B}^T)(\delta_i\mathbf{B}\mathbf{B}^T + \boldsymbol{\Psi})^{-1}(\widetilde{\mathbf{y}}_i - \boldsymbol{\Theta}\widetilde{\mathbf{x}}_i) = \delta_i\mathbf{B}^T(\delta_i\mathbf{B}\mathbf{B}^T + \boldsymbol{\Psi})^{-1}(\widetilde{\mathbf{y}}_i - \boldsymbol{\Theta}\widetilde{\mathbf{x}}_i),$$

and the conditional variance is

$$\text{Var}(\widetilde{\mathbf{z}}_i \mid \widetilde{\mathbf{y}}_i) = \delta_i\mathbf{I}_K - (\delta_i\mathbf{B}^T)(\delta_i\mathbf{B}\mathbf{B}^T + \boldsymbol{\Psi})^{-1}(\delta_i\mathbf{B}) = \delta_i\mathbf{I}_K - \delta_i^2\mathbf{B}^T(\delta_i\mathbf{B}\mathbf{B}^T + \boldsymbol{\Psi})^{-1}\mathbf{B}.$$

To simplify the expressions, let $\mathbf{W}_i = \delta_i\mathbf{B}^T(\delta_i\mathbf{B}\mathbf{B}^T + \boldsymbol{\Psi})^{-1}$ and $\widetilde{\boldsymbol{\epsilon}}_i^* = \widetilde{\mathbf{y}}_i - \boldsymbol{\Theta}\widetilde{\mathbf{x}}_i$.
It follows that, the conditional distribution of $\widetilde{\mathbf{z}}_i|\widetilde{\mathbf{y}}_i \sim \text{MVN}_Q(\mathbf{W}_i\widetilde{\boldsymbol{\epsilon}}_i^*, \delta_i(\mathbf{I}_K - \mathbf{W}_i\mathbf{B}))$.

To calculate the $Q$-function, we can first write

$$
\begin{aligned}
L(\boldsymbol{\Theta},\mathbf{B},\boldsymbol{\Psi}) &\propto \frac{1}{2}\sum_{i=1}^{N}(\widetilde{\boldsymbol{\epsilon}}_i^* - \mathbf{B}\widetilde{\mathbf{z}}_i)^T\boldsymbol{\Psi}^{-1}(\widetilde{\boldsymbol{\epsilon}}_i^* - \mathbf{B}\widetilde{\mathbf{z}}_i) + \frac{n}{2}\log|\boldsymbol{\Psi}| + g_{\lambda_1,\lambda_2}(\boldsymbol{\Theta}) \\
&= \frac{1}{2}\left[\sum_{i=1}^{N}\left(\widetilde{\boldsymbol{\epsilon}}_i^{*T}\boldsymbol{\Psi}^{-1}\widetilde{\boldsymbol{\epsilon}}_i^* - 2\widetilde{\boldsymbol{\epsilon}}_i^{*T}\boldsymbol{\Psi}^{-1}\mathbf{B}\widetilde{\mathbf{z}}_i + \widetilde{\mathbf{z}}_i^T\mathbf{B}^T\boldsymbol{\Psi}^{-1}\mathbf{B}^T\widetilde{\mathbf{z}}_i\right)\right] + \frac{n}{2}\log|\boldsymbol{\Psi}| \\
&+ g_{\lambda_1,\lambda_2}(\boldsymbol{\Theta}) \\
&= \frac{1}{2}\left[\sum_{i=1}^{N}\left\{\widetilde{\boldsymbol{\epsilon}}_i^{*T}\boldsymbol{\Psi}^{-1}\widetilde{\boldsymbol{\epsilon}}_i^* - 2\widetilde{\boldsymbol{\epsilon}}_i^{*T}\boldsymbol{\Psi}^{-1}\mathbf{B}\widetilde{\mathbf{z}}_i + \operatorname{tr}(\mathbf{B}^T\boldsymbol{\Psi}^{-1}\mathbf{B}^T\widetilde{\mathbf{z}}_i\widetilde{\mathbf{z}}_i^T)\right\}\right] + \frac{n}{2}\log|\boldsymbol{\Psi}| \\
&+ g_{\lambda_1,\lambda_2}(\boldsymbol{\Theta})
\end{aligned}
$$

Then, the $Q$-function is given by

$$
\begin{aligned}
Q(\boldsymbol{\Theta},\mathbf{B},\boldsymbol{\Psi}) &\propto \frac{1}{2}\left[\sum_{i=1}^{N}\left\{\widetilde{\boldsymbol{\epsilon}}_i^{*T}\boldsymbol{\Psi}^{-1}\widetilde{\boldsymbol{\epsilon}}_i^* - 2\widetilde{\boldsymbol{\epsilon}}_i^{*T}\boldsymbol{\Psi}^{-1}\mathbf{B}\mathrm{E}(\widetilde{\mathbf{z}}_i \mid \widetilde{\mathbf{y}}_i) + \operatorname{tr}(\mathbf{B}^T\boldsymbol{\Psi}^{-1}\mathbf{B}^T\mathrm{E}(\widetilde{\mathbf{z}}_i\widetilde{\mathbf{z}}_i^T \mid \widetilde{\mathbf{y}}_i))\right\}\right] \\
&+ \frac{n}{2}\log|\boldsymbol{\Psi}| + g_{\lambda_1,\lambda_2}(\boldsymbol{\Theta})
\end{aligned}
$$

It follows that

$$
\begin{aligned}
\mathrm{E}(\widetilde{\mathbf{z}}_i\widetilde{\mathbf{z}}_i^T \mid \widetilde{\mathbf{y}}_i) &= \mathrm{Var}(\widetilde{\mathbf{z}}_i \mid \widetilde{\mathbf{y}}_i) + \mathrm{E}(\widetilde{\mathbf{z}}_i \mid \widetilde{\mathbf{y}}_i)\mathrm{E}(\widetilde{\mathbf{z}}_i \mid \widetilde{\mathbf{y}}_i)^T \\
&= \delta_i(\mathbf{I}_K - \mathbf{W}_i\mathbf{B}) + (\mathbf{W}_i\widetilde{\boldsymbol{\epsilon}}_i^*)(\mathbf{W}_i\widetilde{\boldsymbol{\epsilon}}_i^*)^T \\
&= \delta_i(\mathbf{I}_K - \mathbf{W}_i\mathbf{B}) + \mathbf{W}_i\widetilde{\boldsymbol{\epsilon}}_i^*\widetilde{\boldsymbol{\epsilon}}_i^{*T}\mathbf{W}_i^T
\end{aligned}
$$

In the M-step, we maximize $Q(\boldsymbol{\Theta},\mathbf{B},\boldsymbol{\Psi})$ with respect to $\boldsymbol{\Theta},\mathbf{B}$ and $\boldsymbol{\Psi}$, which will be implemented using the score equations. Let

$$
\begin{aligned}
Q^*(\boldsymbol{\Theta},\mathbf{B},\boldsymbol{\Psi}) &= \frac{1}{2}\left[\sum_{i=1}^{N}\left\{\widetilde{\boldsymbol{\epsilon}}_i^{*T}\boldsymbol{\Psi}^{-1}\widetilde{\boldsymbol{\epsilon}}_i^* - 2\widetilde{\boldsymbol{\epsilon}}_i^{*T}\boldsymbol{\Psi}^{-1}\mathbf{B}\mathrm{E}(\widetilde{\mathbf{z}}_i \mid \widetilde{\mathbf{y}}_i) + \operatorname{tr}(\mathbf{B}^T\boldsymbol{\Psi}^{-1}\mathbf{B}^T\mathrm{E}(\widetilde{\mathbf{z}}_i\widetilde{\mathbf{z}}_i^T \mid \widetilde{\mathbf{y}}_i))\right\}\right] \\
&+ \frac{n}{2}\log|\boldsymbol{\Psi}|.
\end{aligned}
$$

That is, $Q(\boldsymbol{\Theta},\mathbf{B},\boldsymbol{\Psi}) \propto Q^*(\boldsymbol{\Theta},\mathbf{B},\boldsymbol{\Psi}) + g_{\lambda_1,\lambda_2}(\boldsymbol{\Theta})$. Before taking the first-order derivatives, we can rewrite $Q$ function with respect to the parameters of interest. They are

$$
\begin{aligned}
Q(\boldsymbol{\Theta}) \propto Q^*(\boldsymbol{\Theta}) + g_{\lambda_1,\lambda_2}(\boldsymbol{\Theta}) &= -\sum_{i=1}^{N}\widetilde{\mathbf{y}}_i^T\boldsymbol{\Psi}^{-1}\boldsymbol{\Theta}\widetilde{\mathbf{x}}_i - \frac{1}{2}\sum_{i=1}^{N}\widetilde{\mathbf{x}}_i^T\boldsymbol{\Theta}^T\boldsymbol{\Psi}^{-1}\boldsymbol{\Theta}\widetilde{\mathbf{x}}_i \\
&+ \sum_{i=1}^{N}\widetilde{\mathbf{x}}_i^T\boldsymbol{\Theta}^T\boldsymbol{\Psi}^{-1}\mathbf{B}\mathrm{E}(\widetilde{\mathbf{z}}_i \mid \widetilde{\mathbf{y}}_i) + g_{\lambda_1,\lambda_2}(\boldsymbol{\Theta}), \\
Q(\mathbf{B}) \propto Q^*(\mathbf{B}) &= -\sum_{i=1}^{N}\widetilde{\boldsymbol{\epsilon}}_i^{*T}\boldsymbol{\Psi}^{-1}\mathbf{B}\mathrm{E}(\widetilde{\mathbf{z}}_i \mid \widetilde{\mathbf{y}}_i)
\end{aligned}
$$

$$+\frac{1}{2}\sum_{i=1}^{N}\mathrm{tr}(\mathbf{B}^T\mathbf{\Psi}^{-1}\mathbf{B}^T\mathrm{E}(\widetilde{\mathbf{z}}_i\widetilde{\mathbf{z}}_i^T \mid \widetilde{\mathbf{y}}_i)),$$

$$Q(\mathbf{\Psi}) \propto Q^*(\mathbf{\Psi}) = \frac{1}{2}\sum_{i=1}^{N}\widetilde{\boldsymbol{\epsilon}}_i^{*T}\mathbf{\Psi}^{-1}\widetilde{\boldsymbol{\epsilon}}_i^{*} - \sum_{i=1}^{N}\widetilde{\boldsymbol{\epsilon}}_i^{*T}\mathbf{\Psi}^{-1}\mathbf{B}\mathrm{E}(\widetilde{\mathbf{z}}_i \mid \widetilde{\mathbf{y}}_i)$$
$$+\frac{1}{2}\sum_{i=1}^{N}\mathrm{tr}(\mathbf{B}^T\mathbf{\Psi}^{-1}\mathbf{B}^T\mathrm{E}(\widetilde{\mathbf{z}}_i\widetilde{\mathbf{z}}_i^T \mid \widetilde{\mathbf{y}}_i)) + \frac{n}{2}\log|\mathbf{\Psi}|.$$

Let $\widetilde{\mathbf{y}}_i^* = \widetilde{\mathbf{y}}_i - \mathbf{B}\mathrm{E}(\widetilde{\mathbf{z}}_i|\widetilde{\mathbf{y}}_i)$ and $\widetilde{\mathbf{Y}}^*_{N\times Q} = (\widetilde{\mathbf{y}}_1^*, \cdots, \widetilde{\mathbf{y}}_N^*)^T$. Some simple calculations lead to

$$\frac{\partial Q^*(\mathbf{\Theta})}{\partial \mathbf{\Theta}} = -\sum_{i=1}^{N}\mathbf{\Psi}^{-1}\widetilde{\mathbf{y}}_i\widetilde{\mathbf{x}}_i^T + \sum_{i=1}^{N}\mathbf{\Psi}^{-1}\mathbf{\Theta}\widetilde{\mathbf{x}}_i\widetilde{\mathbf{x}}_i^T + \sum_{i=1}^{N}\mathbf{\Psi}^{-1}\mathbf{B}\mathrm{E}(\widetilde{\mathbf{z}}_i \mid \widetilde{\mathbf{y}}_i)\widetilde{\mathbf{x}}_i^T$$
$$= -\sum_{i=1}^{N}\mathbf{\Psi}^{-1}\widetilde{\mathbf{y}}_i^*\widetilde{\mathbf{x}}_i^T + \sum_{i=1}^{N}\mathbf{\Psi}^{-1}\mathbf{\Theta}\widetilde{\mathbf{x}}_i\widetilde{\mathbf{x}}_i^T$$
$$= -\mathbf{\Psi}^{-1}(\widetilde{\mathbf{Y}}^{*T}\widetilde{\mathbf{X}} - \mathbf{\Theta}\widetilde{\mathbf{X}}^T\widetilde{\mathbf{X}})$$

$$\frac{\partial Q(\mathbf{B})}{\partial \mathbf{B}} = \frac{\partial Q^*(\mathbf{B})}{\partial \mathbf{B}} = -\sum_{i=1}^{N}\mathbf{\Psi}^{-1}\widetilde{\boldsymbol{\epsilon}}_i^*\mathrm{E}(\widetilde{\mathbf{z}}_i^T \mid \widetilde{\mathbf{y}}_i) + \sum_{i=1}^{N}\mathbf{\Psi}^{-1}\mathbf{B}\mathrm{E}(\widetilde{\mathbf{z}}_i\widetilde{\mathbf{z}}_i^T \mid \widetilde{\mathbf{y}}_i),$$

$$\frac{\partial Q(\mathbf{\Psi})}{\partial \mathbf{\Psi}^{-1}} = \frac{\partial Q^*(\mathbf{\Psi})}{\partial \mathbf{\Psi}^{-1}} = \frac{1}{2}\sum_{i=1}^{N}\widetilde{\boldsymbol{\epsilon}}_i^*\widetilde{\boldsymbol{\epsilon}}_i^{*T} - \sum_{i=1}^{N}\widetilde{\boldsymbol{\epsilon}}_i^*\mathrm{E}(\widetilde{\mathbf{z}}_i^T \mid \widetilde{\mathbf{y}}_i)\mathbf{B}^T - \sum_{i=1}^{N}\mathbf{B}\mathrm{E}(\widetilde{\mathbf{z}}_i\widetilde{\mathbf{z}}_i^T \mid \widetilde{\mathbf{y}}_i)\mathbf{B}^T - \frac{N}{2}\mathbf{\Psi}$$

Simplifying yields,

$$\frac{\partial Q^*(\mathbf{\Theta})}{\partial \theta_{jk}} = -\psi_i^{-1}\left[\sum_{i=1}^{N} y_{ij}^*\widetilde{x}_{ik} - \sum_{p=1}^{P}\theta_{jp}\sum_{i=1}^{N}\widetilde{x}_{ik}\widetilde{x}_{ip}\right]$$

where $\widetilde{\mathbf{y}}_i^* = (\widetilde{y}_{i1}^*, \ldots, \widetilde{y}_{iQ}^*)^T$ for $i = 1, \ldots, N$.
Notice that

$$\frac{\partial g_{\lambda_1,\lambda_2}(\mathbf{\Theta})}{\partial \theta_{jk}} = \lambda_1\mathrm{sign}(\theta_{jk}) + \lambda_2\frac{\theta_{jk}}{\sqrt{\theta_{1k}^2 + \cdots + \theta_{Qk}^2}}$$
$$= \lambda_1\mathrm{sign}(\theta_{jk}) + \lambda_2\frac{\theta_{jk}}{\|\mathbf{\Theta}_k\|_2},$$

where

$$\mathrm{sign}(x) = \begin{cases} 1 & \text{if } x > 0, \\ 0 & \text{if } x = 0, \\ -1 & \text{if } x < 0, \end{cases}$$

$\mathbf{\Theta}_k$ is the $k$th column of $\mathbf{\Theta}$ and $\|\mathbf{\Theta}_k\|_2 = \sqrt{\theta_{1k}^2 + \cdots + \theta_{Qk}^2}$ denotes the $L_2$ norm of $\mathbf{\Theta}_k$. Setting the above first-order derivatives to zero, we obtain the roots, respectively,

$$\theta_{ij} = \left(1 - \frac{\lambda_2 \psi_i}{a_j \|\boldsymbol{\Theta}_{\text{lasso},j}\|_2}\right) \frac{S(b_{ij}, \lambda_1 \psi_i)}{a_j}, \tag{15}$$

$$\mathbf{B} = \left(\sum_{i=1}^{N} \widetilde{\boldsymbol{\epsilon}}_i^* \mathrm{E}(\widetilde{\mathbf{z}}_i^T \mid \widetilde{\mathbf{y}}_i)\right)\left(\sum_{i=1}^{N} \mathrm{E}(\widetilde{\mathbf{z}}_i \widetilde{\mathbf{z}}_i^T \mid \widetilde{\mathbf{y}}_i)\right)^{-1},$$

$$\boldsymbol{\Psi} = \frac{1}{N} \mathrm{diag}\left(\sum_{i=1}^{N} \widetilde{\boldsymbol{\epsilon}}_i^* \widetilde{\boldsymbol{\epsilon}}_i^{*T} - \sum_{i=1}^{N} \mathbf{B}\mathrm{E}(\widetilde{\mathbf{z}}_i \widetilde{\mathbf{z}}_i^T \mid \widetilde{\mathbf{y}}_i) \mathbf{B}^T\right),$$

where we define a soft-thresholding function

$$S(b, \lambda) = \begin{cases} b - \lambda & \text{if } b > \lambda, \\ b + \lambda & \text{if } b < -\lambda, \\ 0 & \text{otherwise }, \end{cases}$$

$$a_k = \sum_{i=1}^{N} X_{ik}^2 \text{ and } b_{jk} = \sum_{i=1}^{N} X_{ik}(Y_{ij} - \sum_{p \neq k} \theta_{\text{lasso},jp} X_{ip}),$$

and the regularized solution is related to lasso via the following relation [15]:

$$\boldsymbol{\Theta}_k = \frac{a_k}{a_k + \frac{\lambda_2}{\|\boldsymbol{\Theta}_k\|_2}} \boldsymbol{\Theta}_{\text{lasso},k},$$

$\boldsymbol{\Theta}_{\text{lasso},k}$ is the multivariate lasso solution (when $\lambda_2 = 0$). For detailed derivation of the expression to update $\theta_{ij}$, we refer the reader to the Supplementary Material in [15].

### B Derivation of debiased estimator

Following the previous section, the generalized multivariate FARM can be written in a multivariate regression form:

$$\widetilde{\mathbf{y}}_i^* := \widetilde{\mathbf{y}}_i - \mathbf{B}\widetilde{\mathbf{z}}_i = \boldsymbol{\Theta}\widetilde{\mathbf{x}}_i + \widetilde{\boldsymbol{\epsilon}}_i, \quad i = 1, \ldots, N$$

where $\widetilde{\mathbf{Y}}^* = (\widetilde{\mathbf{y}}_1^*, \ldots, \widetilde{\mathbf{y}}_N^*)^T$.

Then, the loss function for the DrFARM problem as a function of $\boldsymbol{\Theta}$ is given by:

$$l(\boldsymbol{\Theta}) \propto -\sum_{i=1}^{N} \widetilde{\mathbf{y}}_i^{*T} \boldsymbol{\Psi}^{-1} \boldsymbol{\Theta}\widetilde{\mathbf{x}}_i + \frac{1}{2} \sum_{i=1}^{N} \widetilde{\mathbf{x}}_i^T \boldsymbol{\Theta}^T \boldsymbol{\Psi}^{-1} \boldsymbol{\Theta}\widetilde{\mathbf{x}}_i + g_{\lambda_1,\lambda_2}(\boldsymbol{\Theta}),$$

where

$$
\begin{aligned}
g_{\lambda_1,\lambda_2}(\boldsymbol{\Theta}) :&= \lambda_1\|\boldsymbol{\Theta}\|_1 + \lambda_2\|\boldsymbol{\Theta}^T\|_{2,1} \\
&= \lambda_1 \sum_{q=1}^{Q}\sum_{p=1}^{P}|\theta_{qp}| + \lambda_2 \sum_{p=1}^{P}\sqrt{\theta_{1p}^2 + \cdots + \theta_{Qp}^2}.
\end{aligned}
$$

To derive the debiased estimator for DrFARM, we take the first derivative

$$
\frac{\partial l(\boldsymbol{\Theta})}{\partial \boldsymbol{\Theta}} = -\boldsymbol{\Psi}^{-1}(\widetilde{\mathbf{Y}}^{*T}\widetilde{\mathbf{X}} - \boldsymbol{\Theta}\widetilde{\mathbf{X}}^T\widetilde{\mathbf{X}}) + \frac{\partial g_{\lambda_1,\lambda_2}(\boldsymbol{\Theta})}{\partial \boldsymbol{\Theta}}.
$$

Setting the derivative zero yields

$$
-\boldsymbol{\Psi}^{-1}(\widetilde{\mathbf{Y}}^{*T}\widetilde{\mathbf{X}} - \boldsymbol{\Theta}\widetilde{\mathbf{X}}^T\widetilde{\mathbf{X}}) + \frac{\partial g_{\lambda_1,\lambda_2}(\boldsymbol{\Theta})}{\partial \boldsymbol{\Theta}} = 0, \tag{16}
$$

and the solution $\widehat{\boldsymbol{\Theta}}$ satisfies the Karush-Kuhn-Tucker (KKT) condition which is given by

$$
\frac{\partial g_{\lambda_1,\lambda_2}(\boldsymbol{\Theta})}{\partial \boldsymbol{\Theta}} = \boldsymbol{\Psi}^{-1}(\widetilde{\mathbf{Y}}^{*T}\widetilde{\mathbf{X}} - \boldsymbol{\Theta}\widetilde{\mathbf{X}}^T\widetilde{\mathbf{X}}). \tag{17}
$$

Recall that $\widehat{\mathbf{C}} = (\mathbf{X}^T\mathbf{X})/N$, rearranging (2) with $\boldsymbol{\Theta} = \widehat{\boldsymbol{\Theta}}$ gives

$$
\widehat{\boldsymbol{\Theta}}\widehat{\mathbf{C}} + \frac{1}{N}\boldsymbol{\Psi}\frac{\partial g_{\lambda_1,\lambda_2}(\boldsymbol{\Theta})}{\partial \boldsymbol{\Theta}}\bigg|_{\boldsymbol{\Theta}=\widehat{\boldsymbol{\Theta}}} = \frac{1}{N}\widetilde{\mathbf{Y}}^{*T}\widetilde{\mathbf{X}}.
$$

Suppose the true parameter is given by $\boldsymbol{\Theta}_0$, deducting $\boldsymbol{\Theta}_0\widehat{\mathbf{C}}$ from the both sides of the equation yields

$$
\begin{aligned}
(\widehat{\boldsymbol{\Theta}} - \boldsymbol{\Theta}_0)\widehat{\mathbf{C}} + \frac{1}{N}\boldsymbol{\Psi}\frac{\partial g_{\lambda_1,\lambda_2}(\boldsymbol{\Theta})}{\partial \boldsymbol{\Theta}}\bigg|_{\boldsymbol{\Theta}=\widehat{\boldsymbol{\Theta}}} &= \frac{1}{N}(\widetilde{\mathbf{Y}}^{*T} - \boldsymbol{\Theta}_0\widetilde{\mathbf{X}}^T)\widetilde{\mathbf{X}} \\
&= \frac{1}{N}\widetilde{\mathbf{E}}^T\widetilde{\mathbf{X}}.
\end{aligned}
$$

Then, we post-multiply the both sides of the equation with $\boldsymbol{\Omega}$ and obtain

$$
\begin{aligned}
&(\widehat{\boldsymbol{\Theta}} - \boldsymbol{\Theta}_0)\widehat{\mathbf{C}}\boldsymbol{\Omega} + \frac{1}{N}\boldsymbol{\Psi}\frac{\partial g_{\lambda_1,\lambda_2}(\boldsymbol{\Theta})}{\partial \boldsymbol{\Theta}}\bigg|_{\boldsymbol{\Theta}=\widehat{\boldsymbol{\Theta}}}\boldsymbol{\Omega} = \frac{1}{N}\widetilde{\mathbf{E}}^T\widetilde{\mathbf{X}}\boldsymbol{\Omega} \\
\implies &(\widehat{\boldsymbol{\Theta}} - \boldsymbol{\Theta}_0) + \frac{1}{N}\boldsymbol{\Psi}\frac{\partial g_{\lambda_1,\lambda_2}(\boldsymbol{\Theta})}{\partial \boldsymbol{\Theta}}\bigg|_{\boldsymbol{\Theta}=\widehat{\boldsymbol{\Theta}}}\boldsymbol{\Omega} = \frac{1}{N}\widetilde{\mathbf{E}}^T\widetilde{\mathbf{X}}\boldsymbol{\Omega} - (\widehat{\boldsymbol{\Theta}} - \boldsymbol{\Theta}_0)(\widehat{\mathbf{C}}\boldsymbol{\Omega} - \mathbf{I}_p) \\
\implies &(\widehat{\boldsymbol{\Theta}} - \boldsymbol{\Theta}_0) + \frac{1}{N}\boldsymbol{\Psi}\frac{\partial g_{\lambda_1,\lambda_2}(\boldsymbol{\Theta})}{\partial \boldsymbol{\Theta}}\bigg|_{\boldsymbol{\Theta}=\widehat{\boldsymbol{\Theta}}}\boldsymbol{\Omega} = \frac{1}{N}\widetilde{\mathbf{E}}^T\widetilde{\mathbf{X}}\boldsymbol{\Omega} - \frac{\boldsymbol{\Delta}}{\sqrt{N}},
\end{aligned}
$$

where $\boldsymbol{\Delta} = \sqrt{N}(\widehat{\boldsymbol{\Theta}} - \boldsymbol{\Theta}_0)(\boldsymbol{\Omega}\widehat{\mathbf{C}} - \mathbf{I}_p)$.
Let $W_i = \frac{1}{\sqrt{N}}\widetilde{\mathbf{E}}_i^T\mathbf{X}\boldsymbol{\Omega}$. Since $\widetilde{\mathbf{E}} \sim \mathrm{MN}_{N\times Q}(\mathbf{O}_{N\times Q}, \mathbf{I}_N, \boldsymbol{\Psi})$, we have $\widetilde{\mathbf{E}}_i \sim \mathrm{MVN}_N(\mathbf{0}_N, \psi_i\mathbf{I}_N)$ and

$$\mathrm{Var}(W_i^T) = \frac{1}{N}\boldsymbol{\Omega}\widetilde{\mathbf{X}}^T\psi_i\mathbf{I}_N\widetilde{\mathbf{X}}\boldsymbol{\Omega}^T = \psi_i\boldsymbol{\Omega}\widehat{\mathbf{C}}\boldsymbol{\Omega}^T.$$

It follows that $W_i^T \sim \mathrm{MVN}_N(\mathbf{0}_N, \psi_i\boldsymbol{\Phi})$, where $\boldsymbol{\Phi} := \boldsymbol{\Omega}\widehat{\mathbf{C}}\boldsymbol{\Omega}^T$.
Suppose that $\|\boldsymbol{\Delta}^T\|_\infty = \max_i \|\boldsymbol{\Delta}_i^T\|_\infty = o_p(1)$ (see [21, 23]) where

$$\boldsymbol{\Delta} := \left[\boldsymbol{\Delta}_1^T \ \boldsymbol{\Delta}_2^T \ \ldots \ \boldsymbol{\Delta}_Q^T\right]^T,$$

then

$$\sqrt{N}\left(\left(\widehat{\boldsymbol{\Theta}} + \frac{1}{N}\boldsymbol{\Psi}\frac{\partial g_{\lambda_1,\lambda_2}(\boldsymbol{\Theta})}{\partial\boldsymbol{\Theta}}\bigg|_{\boldsymbol{\Theta}=\widehat{\boldsymbol{\Theta}}}\right) - \boldsymbol{\Theta}_0\right) = \left[W_1^T \ W_2^T \ \ldots \ W_Q^T\right]^T + \left[o_p(1) \ o_p(1) \ \ldots \ o_p(1)\right]^T,$$

and we can define the debiased estimator for DrFARM based on

$$\widehat{\boldsymbol{\Theta}}^*_{\mathrm{db}} = \widehat{\boldsymbol{\Theta}} + \frac{1}{N}\boldsymbol{\Psi}\frac{\partial g_{\lambda_1,\lambda_2}(\boldsymbol{\Theta})}{\partial\boldsymbol{\Theta}}\bigg|_{\boldsymbol{\Theta}=\widehat{\boldsymbol{\Theta}}}\boldsymbol{\Omega} = \widetilde{\boldsymbol{\Theta}} = \widehat{\boldsymbol{\Theta}} + \frac{1}{N}(\widetilde{\mathbf{Y}}^{*T} - \widehat{\boldsymbol{\Theta}}\widetilde{\mathbf{X}}^T)\widetilde{\mathbf{X}}\boldsymbol{\Omega},$$

where $\widehat{\boldsymbol{\Theta}}^*_{\mathrm{db}} = \{\hat{\theta}^*_{\mathrm{db},ij}\}$ and

$$\frac{\sqrt{N}(\hat{\theta}^*_{\mathrm{db},ij} - \theta_{ij})}{\sqrt{\psi_i\boldsymbol{\Phi}_{jj}}} \xrightarrow{d} \mathrm{N}(0,1), \ \text{as } N \to \infty$$

.
Notice that $\widehat{\boldsymbol{\Theta}}^*_{\mathrm{db}}$ takes resemblance to the lasso counterpart:

$$\widehat{\boldsymbol{\beta}}_{\mathrm{db}} = \widehat{\boldsymbol{\beta}} + \frac{1}{n}\widehat{\boldsymbol{\Omega}}\mathbf{X}^T(Y - \mathbf{X}\widehat{\boldsymbol{\beta}}).$$

However, since we do not observe $\widetilde{\mathbf{Z}}$ in $\widetilde{\mathbf{Y}}^*$, we replace $\widetilde{\mathbf{Z}}$ by $\widehat{\mathrm{E}}(\widetilde{\mathbf{Z}}|\widetilde{\mathbf{Y}})$ and using a working version of $\widetilde{\mathbf{Y}}^*$ given by $\widetilde{\mathbf{Y}} - \widehat{\mathrm{E}}(\widetilde{\mathbf{Z}}|\widetilde{\mathbf{Y}})\widehat{\mathbf{B}}^T$. This gives rise to the following estimator:

$$\widehat{\boldsymbol{\Theta}}_{\mathrm{db}} = \widehat{\boldsymbol{\Theta}} + \frac{1}{N}(\widetilde{\mathbf{Y}}^T - \widehat{\boldsymbol{\Theta}}\widetilde{\mathbf{X}}^T - \widehat{\mathbf{B}}\widehat{\mathrm{E}}(\widetilde{\mathbf{Z}} \mid \widetilde{\mathbf{Y}})^T)\widetilde{\mathbf{X}}\widehat{\boldsymbol{\Omega}}.$$

### C Implementation

We normalize all the columns of the predictors and traits to have a mean zero and variance one so that they are on the same scale prior to the operation of regularization. Let the normalized predictor matrix and normalized trait matrices be $\mathbf{X}$ and $\mathbf{Y}$, respectively. We denote the de-association transformed predictors and trait matrices by $\widetilde{\mathbf{X}}$ and $\widetilde{\mathbf{Y}}$ (note that $\widetilde{\mathbf{X}} = \mathbf{X}$

and $\widetilde{\mathbf{Y}} = \mathbf{Y}$ in simulation 1 and carry out calculations given in Algorithm 1. The default tolerance $\xi$ is set at $10^{-4}$. We used remMap and searched for the solution against a $10 \times 10$ tuning parameter grid for $(\lambda_1, \lambda_2)$ using the R package `remMap`. Parameter tuning is done with EBIC with $\gamma = 1$ [57] (default). We used the optional remMap solution as is done with the initial value $\mathbf{\Theta}^{(0)}$. With the initial sparse estimate $\mathbf{\Theta}^{(0)}$, we obtained the initial debiased estimate $\mathbf{\Theta}_{\text{db}}^{(0)}$ by equation (5), followed by the initial residual matrix $\widetilde{\mathbf{E}}^{*(0)} = \widetilde{\mathbf{Y}} - \widetilde{\mathbf{X}}\mathbf{\Theta}_{\text{db}}^{(0)}$, which was then analyzed by `fa()` from the R package `psych`, where the maximum likelihood rotation estimation with no rotation is coded to obtain the initial loading estimate $\mathbf{B}^{(0)}$ and the initial uniqueness parameter estimates $\mathbf{\Psi}^{(0)} = \text{diag}(\psi_1^{(0)}, \ldots, \psi_Q^{(0)})$. Suppose $(\lambda_1^*, \lambda_2^*)$ is the optimal tuning parameter selected. Since the scale of tuning parameters $(\lambda_1, \lambda_2)$ are different between remMap and DrFARM, we calculated the minimum $(\psi_{\min})$, first quartile $(\psi_{Q_1})$, median $(\psi_{\text{med}})$, third quartile$(\psi_{Q_3})$ and maximum $(\psi_{\max})$ of the initial uniqueness $\psi_i^{(0)}$'s and searched against the $5 \times 5$ grid $[\lambda_1^*/\psi_{\min}, \lambda_1^*/\psi_{Q_1}, \lambda_1^*/\psi_{\text{med}}, \lambda_1^*/\psi_{Q_3}, \lambda_1^*/\psi_{\max}] \times [\lambda_2^*/\psi_{\min}, \lambda_2^*/\psi_{Q_1}, \lambda_2^*/\psi_{\text{med}}, \lambda_2^*/\psi_{Q_3}, \lambda_2^*/\psi_{\max}]$. The tuning is done by EBIC with $\gamma = 1$.

For simulation 2 (where data was generated using kinship), we used univariate trait lasso to generate $\widehat{\mathbf{\Theta}}^{(0)}$, where we fit $Q$ separate univariate trait lasso regressions $Y_i \sim \mathbf{X}, i = 1, \ldots, Q$ and obtained the tuning parameter $\omega_i$ using 10-fold cross validation with the R package `glmnet`. The coefficients corresponding to the $\omega_i$ are used as the $i$th row of $\widehat{\mathbf{\Theta}}^{(0)}$. Similar to Simulation 1, we searched against a $5 \times 5$ grid $[\omega/\psi_{\min}, \omega/\psi_{Q_1}, \omega/\psi_{\text{med}}, \omega/\psi_{Q_3}, \omega/\psi_{\max}] \times [\omega/\psi_{\min}, \omega/\psi_{Q_1}, \omega/\psi_{\text{med}}, \omega/\psi_{Q_3}, \omega/\psi_{\max}]$, with $\omega = \sum_{i=1}^{Q} \omega_i$.

In general, we observed that remMap provided better initial estimate compared to the method of row-wise lasso, leading to better performances on the inference for $\mathbf{\Theta}$. Therefore, we recommend using the row-wise lasso method only if remMap fails to provide reasonable initial values. We assume the number of latent factors $K$ to be known throughout the simulation studies.

### D Additional Tables

**Table 3** Performance metrics across 100 replicates for remMap ($r$) and DrFARM ($d$) under different type of debiasing in Scenario I for simulation 1. The true negative rate (TNR) and true positive rate (TPR) were not shown for the individual level and group level results, respectively, as all methods achieve close to 100%. NL: Nodewise lasso; QO: Quadratic optimization.

| | | | Individual | | | Group | | |
|---|---|---|---|---|---|---|---|---|
| Method | Precision | Debiasing | TPR | FDR | MCC | TNR | FDR | MCC |
| $d$ | None | None | 94.9% | 17.1% | 88.2% | 88.2% | 35.7% | 75.2% |
| $d$ | Glasso | Outer | 92.2% | 5.2% | 93.5% | 99.5% | 2.5% | 98.5% |
| $d$ | Glasso | Inner | 93.4% | 15.9% | 88.0% | 82.6% | 40.4% | 69.8% |
| $d$ | Glasso | Double | 90.3% | 5.1% | 92.5% | 99.5% | 2.6% | 98.4% |
| $d$ | NL | Inner | 89.2% | 6.2% | 91.3% | 93.8% | 22.5% | 85.1% |
| $d$ | NL | Double | 87.4% | 4.5% | 91.3% | 99.7% | 1.8% | 98.8% |
| $d$ | QO | Inner | 92.6% | 8.7% | 91.8% | 91.4% | 29.3% | 80.1% |
| $d$ | QO | Double | 84.2% | 6.9% | 88.5% | 99.5% | 2.6% | 98.3% |
| $r$ | None | None | 84.8% | 32.2% | 75.1% | 86.2% | 40.2% | 71.4% |
| $r$ | Glasso | Outer | 70.3% | 6.3% | 81.1% | 99.4% | 3.4% | 97.5% |

**Table 4** Performance metrics across 100 replicates for debiased DrFARM under different algorithm in Scenario I for simulation 2. The true negative rate (TNR) true positive (TPR) were not shown for the individual level and group level results as all methods performed similarly ($\sim$ 100% and $\sim$ 99.5%, respectively).

| | Individual | | | Group | | |
|---|---|---|---|---|---|---|
| Algorithm | TPR | FDR | MCC | TPR | FDR | MCC |
| Naïve with $\mathbf{I}_N$ | 85.1% | 5.2% | 89.8% | 96.6% | 1.9% | 96.9% |
| Naïve with $\mathbf{K}$ | 85.0% | 5.2% | 89.8% | 96.6% | 2.1% | 96.8% |
| Glasso with $\mathbf{I}_N$ | 82.1% | 5.2% | 88.2% | 95.7% | 2.2% | 96.1% |
| Glasso with $\mathbf{K}$ | 81.7% | 5.1% | 88.0% | 95.4% | 2.1% | 96.1% |
| NL with $\mathbf{I}_N$ | 76.6% | 4.6% | 85.4% | 93.6% | 1.9% | 95.1% |
| NL with $\mathbf{K}$ | 77.7% | 4.6% | 86.1% | 94.0% | 2.0% | 95.3% |
| QO with $\mathbf{I}_N$ | 74.9% | 7.2% | 83.4% | 93.3% | 2.8% | 94.4% |
| QO with $\mathbf{K}$ | 75.0% | 6.9% | 83.5% | 93.2% | 2.8% | 94.3% |

**Table 5** Performance metrics across 100 replicates for debiased DrFARM under different algorithm in Scenario II for simulation 2. The true negative rate (TNR) were not shown for the individual level results as all methods achieve close to 100%.

| | Individual | | | Group | | | |
|---|---|---|---|---|---|---|---|
| Algorithm | TPR | FDR | MCC | TPR | TNR | FDR | MCC |
| Naïve with $\mathbf{I}_N$ | 93.9% | 5.0% | 94.4% | 99.6% | 99.3% | 4.0% | 97.4% |
| Naïve with $\mathbf{K}$ | 93.9% | 5.0% | 94.4% | 99.6% | 99.3% | 3.9% | 97.4% |
| Glasso with $\mathbf{I}_N$ | 90.8% | 4.6% | 93.0% | 99.0% | 99.3% | 4.0% | 97.0% |
| Glasso with $\mathbf{K}$ | 87.5% | 5.1% | 91.1% | 98.3% | 99.2% | 4.3% | 96.4% |
| NL with $\mathbf{I}_N$ | 90.1% | 4.6% | 92.7% | 98.9% | 99.3% | 3.9% | 97.1% |
| NL with $\mathbf{K}$ | 87.3% | 4.8% | 91.1% | 98.4% | 99.3% | 4.1% | 96.6% |
| QO with $\mathbf{I}_N$ | 86.3% | 9.3% | 88.5% | 98.1% | 98.7% | 7.2% | 94.6% |
| QO with $\mathbf{K}$ | 86.3% | 9.2% | 88.5% | 98.1% | 98.6% | 7.3% | 94.5% |
